# Human iPSC-derived Committed Cardiac Progenitors Generate Cardiac Tissue Grafts in a Swine Ischemic Cardiomyopathy Model without Triggering Ventricular Arrhythmias

**DOI:** 10.1101/2024.02.14.580375

**Authors:** Amish N. Raval, Eric G. Schmuck, Vladislav Leonov, Sushmita Roy, Yukihiro Saito, Tianhua Zhou, Hae Jae Jang, Kaylee Wilker, James Conklin, Timothy A. Hacker, Chad Koonce, Meghan Boyer, Kristin Stack, Ellen Hebron, Scott K. Nagle, Jianhua Zhang, Patrick C.H. Hsieh, Timothy J. Kamp

## Abstract

**Background:** Intramyocardial injection of human pluripotent stem cell-derived cardiomyocytes following a myocardial infarction (MI) improves cardiac function in large animal models, but associated ventricular arrhythmias are major safety concern. We hypothesized that transendocardial injection of human induced pluripotent stem cell (hiPSC)-derived committed cardiac progenitor cells (CCPs), combined with cardiac fibroblast-derived extracellular matrix (cECM) to enhance cell retention, will generate cardiac tissue grafts improving contractility without triggering ventricular arrhythmias.

**Methods:** hiPSCs were differentiated using bioreactors and small molecules to produce committed cardiac progenitor cells (CCPs). MI was created using a coronary artery balloon occlusion and reperfusion model in Yucatan mini pigs. Four weeks later, epicardial needle injections of CCPs+cECM were performed in a small initial feasibility cohort (n=6), and then transendocardial injections of CCPs+cECM (n=14), CCPs alone (n=14), cECM alone (n=4) or vehicle control (n=13) into the peri-infarct region in a randomized cohort. Arrhythmias were evaluated using implanted event recorders. Magnetic resonance imaging (MRI) and invasive pressure-volume assessment were used to evaluate left ventricular anatomic and functional performance. Detailed histology was performed to detect and characterize human grafts.

**Results:** A scalable biomanufacturing protocol was developed generating CCPs which can efficiently differentiate into cardiomyocytes or endothelial cells in vitro. Intramyocardial delivery of CCPs to post-MI porcine hearts resulted in engraftment and differentiation of CCPs to form ventricular cardiomyocyte rich grafts. There was no significant difference in cardiac MRI-based measured cardiac volumes or function between control, CCP and CCP+cECM groups; however, pressure-volume analysis showed an improvement in dobutamine-stimulated functional reserve in CCP and CCP+cECM groups. Delivery of CCPs did not result in tumors or ventricular arrhythmias.

**Conclusions:** Transendocardial delivery of CCPs with or without cECM into post-MI porcine hearts resulted in comparable human cardiomyocyte grafts which did not improve resting LV function but did improve stress-induced contractile reserve without triggering ventricular arrhythmias.

## INTRODUCTION

Currently approved medication and device therapies can blunt the progression of heart failure (HF) following myocardial infarction (MI), but the morbidity and mortality in post – MI survivors remain significant.^1,2^ For patients who suffer large MIs, replacement of necrotic myocardium with fibrotic scar frequently leads to progressive heart failure, and there are currently no available treatments to replace lost cardiomyocytes and restore contractility. Therefore, cell-based therapies have been investigated for their potential to generate functional cardiac tissue in post-MI or ischemic cardiomyopathy animal models and patients. More than two decades of investigations and clinical trials with a variety of adult cell sources such as bone marrow mononuclear cells have produced no clear evidence of new functional heart muscle or consistent improvements in clinical outcomes.^3^ In contrast, post-MI delivery of human pluripotent stem cell (PSC)-derived cardiomyocytes (CMs) from embryonic stem cells (ESCs) or induced pluripotent stem cells (iPSCs) has produced viable cardiomyocyte grafts in rat,^4–7^ guinea pig,^8–10^ swine,^11–14^ and non-human primate^15–17^ myocardium. However, large animal experimental studies using hPSC-CMs have been complicated by ventricular arrhythmias, relatively low cell retention in the myocardium and mixed functional results.

hPSC-derived cardiac progenitor cells are also being investigated for cardiac cell therapy in the post-MI failing heart,^18,19^ but progenitor cells are a less studied cell preparation for therapy relative to hPSC-CMs. However, there are several features that make cardiac progenitors cells a promising cell source. Cardiac progenitor cells are smaller cells than hPSC-CMs and may be more amenable to manipulation and cell delivery while maintaining viability. The hPSC-derived cardiac progenitor cells are proliferative and therefore may generate larger hPSC-CM grafts per cell delivered. Importantly, cardiac progenitor cells can differentiate to not only hPSC-CMs but also other cardiac cell lineages such as endothelial cells and smooth muscle cells needed to generate viable cardiac tissue grafts. The first in human heart failure treatment clinical trial using hPSCs tested early stage hESC-derived progenitors delivered in a fibrin patch at the time of coronary artery bypass surgery and demonstrated safety for this preparation.^18^ However, hPSCs undergo a dynamic range of cell state transitions in the process of differentiating to terminal cardiac lineages which are variably referred to as cardiac progenitors, and the specific characteristics of the given progenitor state chosen for cell therapy is critically important. The optimal cardiac progenitor cell population isolated from differentiating hPSCs and delivery strategy for cardiac repair remain to been determined.

Low cell retention is a vexing problem for all cardiac cell therapies. Our group previously described a cardiac extracellular matrix (cECM) engineered biomaterial using high-density culture-expanded cardiac fibroblasts isolated from human cadaveric hearts. cECM is an acellular, uniquely fibronectin abundant, non-chemically cross-linked, non-hydrogel scaffold with immunomodulatory and angiogenic properties that can be formulated into particulates, reconstituted into solution, and administered as an injectable^20,21^. cECM was shown to educate monocytes to a distinct population of macrophages that exhibit anti-inflammatory and pro-angiogenic features,^21^ and can promote myocardial retention of mesenchymal stem cells in rodent heart disease models when co-delivered epicardially via an open chest.^22,23^ In comparison, a more clinically appealing delivery approach is transendocardial injection (TEI). TEI uses a steerable, needle tipped catheter that is introduced into the left ventricle via a femoral artery to perform targeted intramuscular injections from the endocardial surface.

In this study, we tested whether human iPSC-derived committed cardiac progenitors (CCPs) alone or with cECM could be safely delivered to post-MI failing porcine heart and engraft to form cardiac tissue grafts that lead to improvements in functional outcomes. We observed that the CCPs, regardless of the presence of cECM, successfully engrafted and differentiated into cardiomyocytes, endothelial cells and fibroblasts, leading to the formation of cardiac tissue grafts. Cardiac MRI did not show significant improvement in chamber size or ejection fraction, but pressure-volume analysis demonstrated an improvement in dobutamine-stimulated functional reserve by CCPs. Cell delivery did not induce ventricular arrhythmias in the engrafted hearts.

## METHODS

For detailed methods, please see the Supplemental Methods section in the Supplemental Material. Data and study materials are available from the corresponding authors upon reasonable request. All animal use in this study adhered to an approved protocol by the institutional animal care and use committee at the University of Wisconsin – Madison, protocol, M005512-R03-A01.

### Statistical Analysis

Statistical analysis was carried out using GraphPad Prism version 9. MRI data was analyzed with a 2-way ANOVA with a Sidak post-hoc analysis for multiple comparisons. Pressure volume loops were analyzed with a One-Way ANOVA with a Bonferroni post-hoc analysis. All data presented as mean +/− standard error of the mean unless otherwise stated as standard deviation. Statistical significance was defined as P<0.05.

## RESULTS

### Production of human iPSC-derived CCPs

For scalable production of human iPSC-derived CCPs, iPSCs were propagated as monolayer cultures in CELLSTACK culture chambers. Cells were singularized and inoculated into PBS3 Bioreactors to form cellular aggregates that were differentiated in suspension culture using an accelerated small molecule-based protocol employing biphasic modulation of Wnt signaling along with application of a rho-kinase inhibitor and TGF-β receptor inhibitor (**Figure 1A**). Flow cytometry analysis revealed a progressive loss of expression of the epithelial marker, EpCAM, present on iPSCs and upregulation of the mesodermal marker CD56 (NCAM) during the differentiation protocol (**Figure 1B**).^24^ A KDR^+^/PDGFRα^+^ population emerges and peaks at day 4 consistent with the generation of a cardiac progenitor population.^25,26^ KDR expression subsequently decreases on day 5 and 6 suggesting advancing maturation of these cardiac progenitors. CCPs are identified as the CD56^+^/PDGFRα^+^ or CD56^+^/CXCR4^−^ population present on day 6. Average flow cytometry time course data from 20 runs demonstrate the reproducibility of the differentiation protocol (**Figure 1B, Supplemental Tables 1-4)**. Analysis of gene expression by quantitative RT-PCR over the 6-day protocol revealed loss of expression pluripotency genes (*NANOG, SOX2, POU5F1)* and an increase in expression of cardiac progenitor genes (*HAND2, GATA4, NKX2-5, PDGFRA, TBX5*) (**Figure 1C**). At day 6, the aggregates are dissociated, cryopreserved and described as CCPs.

**Figure 1.**
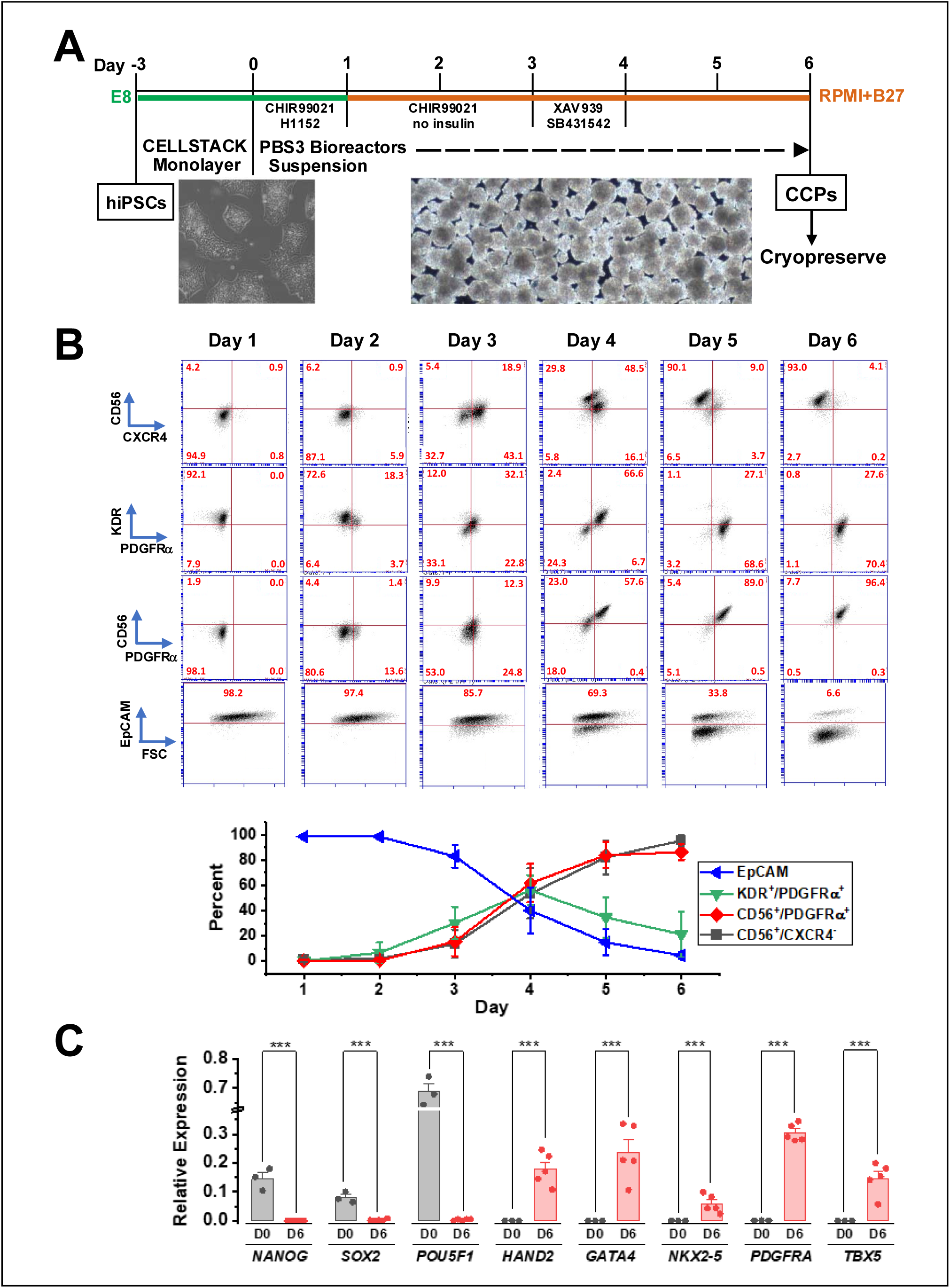
Differentiation of human iPSCs to CCPs. **A)** Differentiation schematic to manufacture CCPs with images showing starting aggregates of human iPSCs and final aggregates of CCPs. **B)** Time course of differentation characterizing cell surface protein expression of CD56, KDR, PDGFRα, and EpCAM using flow cytometry from a representative experiment with average data from 7-20 different runs, error bars ±S.D. **C)** Gene expression of pluripotent and cardiac-specific genes at day 0 (iPSCs) and day 6 (CCPs) relative to the ribosomal protein *RPL32*. Error bars ±S.E.M., ***p<0.005 by t-test comparison.

Single cell RNA-sequencing (scRNA-seq) was also performed to better characterize the CCPs. Cryopreserved CCPs were recovered, and a total of 37,014 cells were analyzed using the 10X Genomics Next GEM single cell 3’ assay. Chromium single cell data output from the 10X Genomics analysis pipeline were further analyzed using Seurat and Cellxgene VIP software. Transcriptomic analyses of the CCPs identified 7 clusters, and 5 clusters were closely related (C0, C1, C2, C4, C6) (**Figure 2A)**. Analysis of differentially expressed genes (DEGs) revealed cardiomyocyte-fated cells in clusters C0, C2, C4 and C6 with expression of genes encoding cardiac transcription factors (*MEF2C, TBX5, NKX2-5*), myofilaments (*ACTC1, MYL4, MYL7, MYH6*), EC coupling proteins (*RYR2*) and ion channels (*CACNA1D, KCNIP4, SLC8A1*). These 4 cell populations account for 64% of the total cells in the CCPs and were annotated cardiomyocyte progenitor (CMP) (**Figure 2B,F**). Further analysis of the DEGs and the GO terms showed the C2 and C4 cell populations differentially express cell cycle marker genes *MKI67* and *TOP2A*. In addition, C2 cells differentially express *CCNB1* (Cyclin B1) and *CENPA* in contrast to C4 cells which express *CDC6* and *FEN1* that are essential for DNA replication, indicating capture of M phase and S phase proliferating CMPs, respectively (**Figure 2E,F, Supplement 2**). The small cluster C6 has abundant expression of *TTN* and *RYR2*, suggesting a differentiating cardiomyocyte (CM). Interestingly, cluster C1 that accounts for 15% of the total CCPs has elevated expression of ribosomal genes including *RPLP1*, *RPS15* as well as genes for ATP synthase complex (*ATP5F1E*) and NADH dehydrogenase (*NDUFS5*) in mitochondrial that are essential for protein translation and metabolism along with lower expression of myofilament, EC coupling and ion channels genes than the CMP populations, so we annotated this cell population as a cardiac progenitor (CP) (**Figure 2A,F**). The C3 and C5 clusters account for 12% and 8.6% of the total CCPs, respectively, and have distinct gene expression pattern from the CMP. The C3 cells differentially express typical endodermal marker genes of EPCAM and FOXA2, and C5 cells differentially express typical endothelial marker genes of CD34, CDH5 (CD144), and PECAM1 (CD31) (**Figure 2C,D,F)**. Overall, the scRNA-seq analysis confirmed that the CCP preparation is composed primarily of cells fated to become CMs (∼80%) whereas a smaller populations of endothelial and endodermal cells are present.

**Figure 2.**
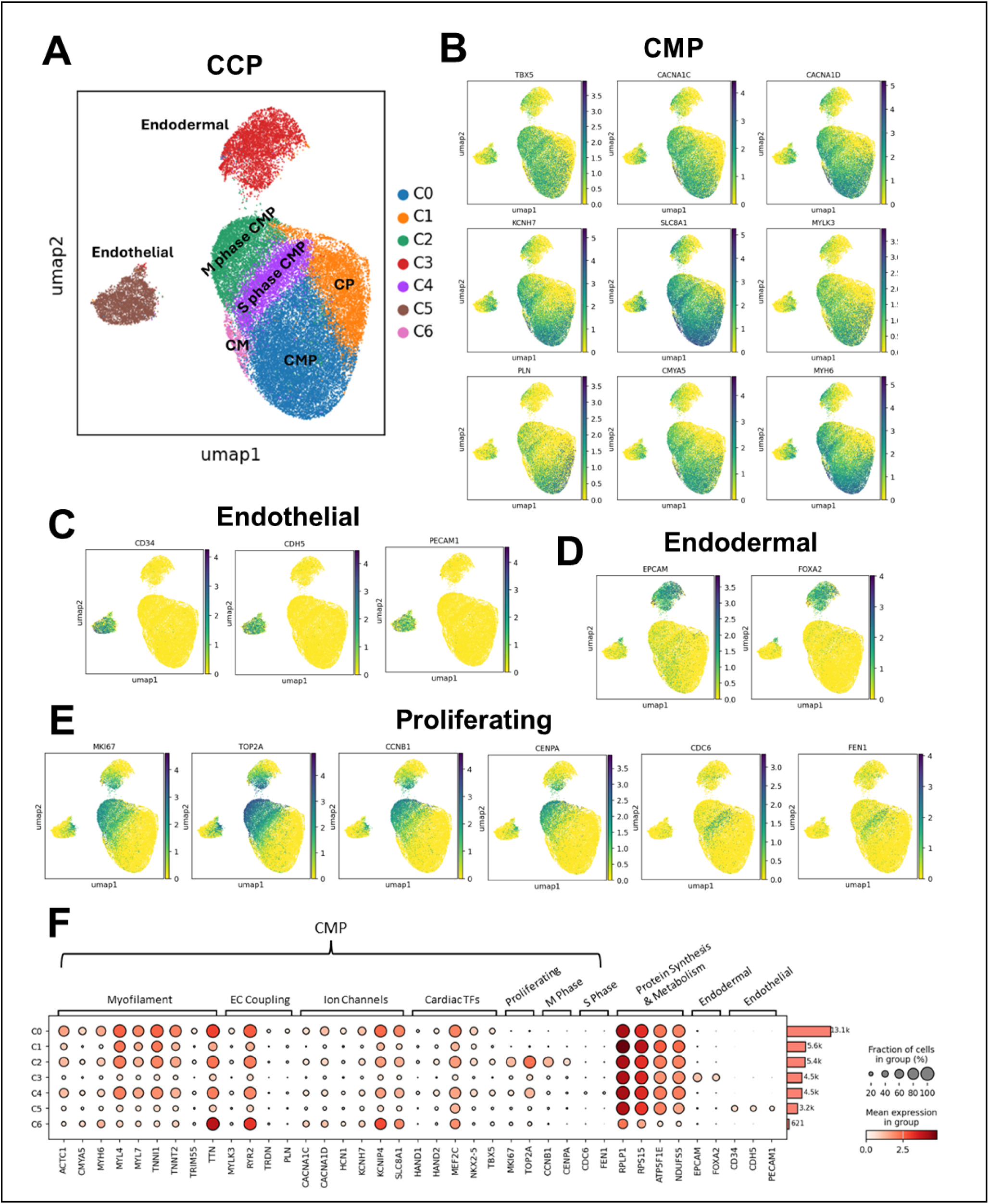
Single cell RNA-sequencing analysis of CCPs. **A)** UMAP projection for CCPs identifies 7 cell clusters annotated based on differential gene expression. **B-E)** Embedding plots for expression of cardiac marker genes in CMP, endothelial cell, endodermal cell, and proliferating cell, respectively. Scale is log2 fold change. **F)** Dot plot showing expression of representative marker genes used to annotate the clusters. CMP: cardiomyocyte progenitor; CP: cardiac progenitor; CM: cardiomyocyte.

### *In vitro* potency of CCPs

To test the potency of the cryopreserved CCPs, cells were thawed and plated on vitronectin substrate in RPMI+B27 medium and characterized after 7 days of culture. After 7 days in culture in RPMI+B27, the cells differentiated into contracting sheets of iPSC-derived cardiomyocytes (iPSC-CMs) (Video supplement 1). Flow cytometry demonstrated that upon thawing 9.3±4.0% of the cells expressed low levels of sarcomeric α-actinin (SAA), but after 7 days of culture 95.9±0.8% of the differentiated cells expressed SAA consistent with differentiation primarily to iPSC-CMs (**Figure 3A**). Immunolabeling of the differentiated cells confirmed that nearly all cells were iPSC-CMs which expressed Nxk2.5, SAA, cardiac troponin I (cTnI), and cardiac troponin T (cTnT) (**Figure 3B**). Optical mapping of the contracting monolayers of iPSC-CMs loaded with a voltage-sensitive fluorescent dye, FluoVolt^TM^, demonstrated optical action potentials with an average spontaneous rate of 1.16±0.02 Hz (n=4). Point stimulation of the cultures over a range of frequencies (1.5-3.0 Hz) showed electrical coupling between the iPSC-CMs with uniform conduction through the monolayers and conduction velocities (CV) ranging from 9.2±0.2 to 13.4±0.4 cm/s (**Figure 3C,D)**. The optical action potential durations progressively shortened at higher rates of stimulation consistent with rate adaptation of cardiac action potentials (**Figure 3C,D).** Single iPSC-CMs (n=27) were characterized with perforated patch clamp recording and demonstrated action potentials when stimulated at 1 Hz with an average maximum diastolic potential (MDP) of –72.5±1.1 mV, action potential amplitude (APA) of 107.0±1.7 mV and dV/dt_max_ of 112.1±14.5 V/s **(Figure 3E, Supplemental Table 5)**. These results show that thawed CCPs plated in basal RPMI+B27 medium without additional growth factors or small molecules rapidly differentiate predominantly to iPSC-CMs and exhibit electrophysiological properties typical of early developing human cardiomyocytes.

**Figure 3.**
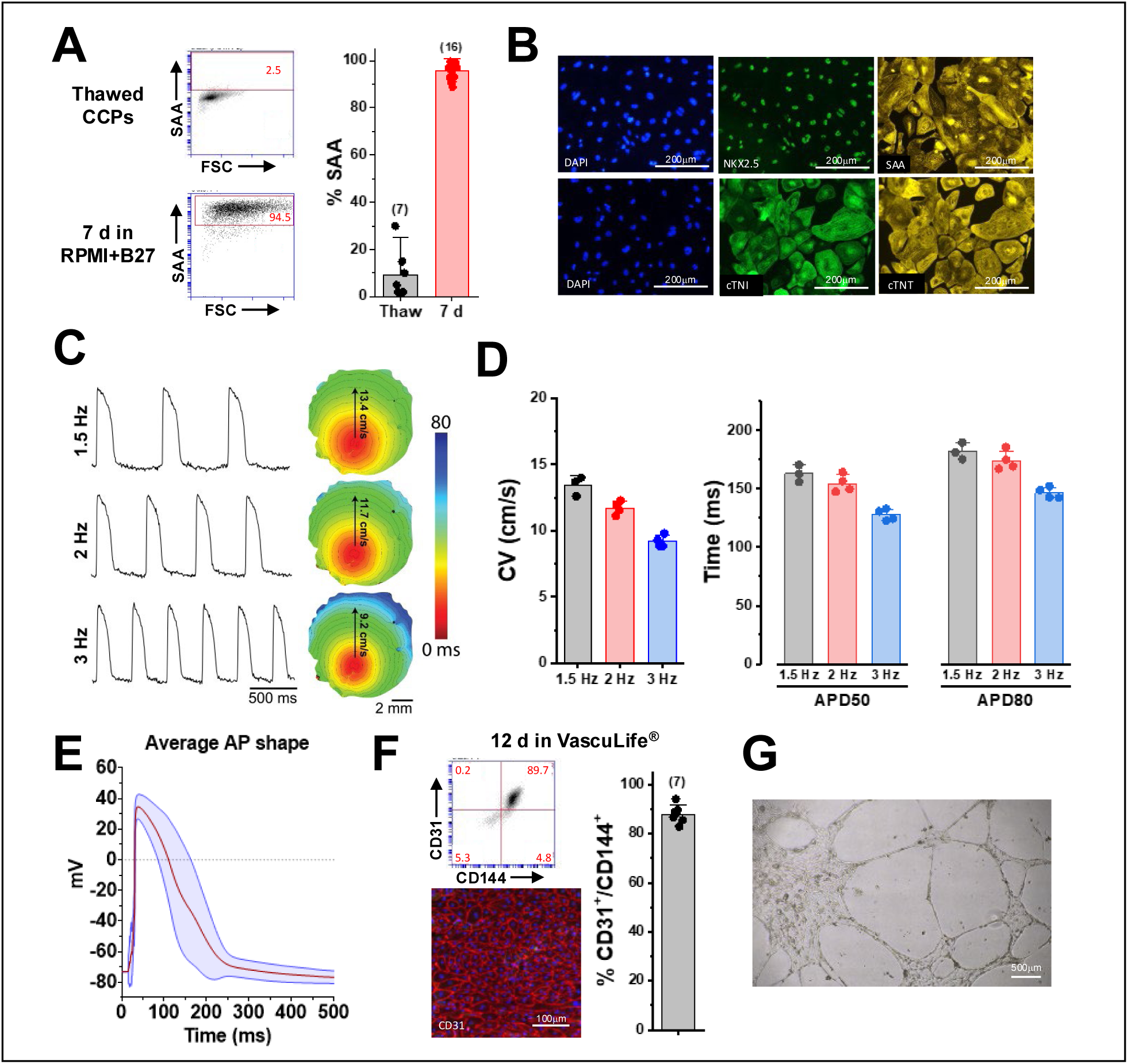
In vitro differentiation of CCPs to iPSC-CMs and iPSC-ECs. **A)** Flow cytometry analysis of CCPs immediately after thaw and following 7 d of culture on vitronectin in RPMI+B27 for sarcomeric α-actinin (SAA) expression. **B)** Immunolabeling of differentiated CCPs after 7 d of culture in RPMI+B27 for cardiomyoctye markers Nkx2.5, SAA, cardiac troponin I (cTnI) and cardiac troponin T (cTnT). **C)** Optical action potentials measured at 1.5, 2.0, and 3.0 Hz stimulation from differentiated iPSC-CM monolayers loaded with Fluovolt^TM^ with conduction maps shown to right. **D)** Average data from optical action potentials for conduction velocity (CV) and action potential duration at 50% (APD50) and 80% (APD80) of repolarization. **E)** Average perforated patch clamp recorded action potential in red and SD in blue for 27 iPSC-CMs. **F)** Flow cytometry for endothelial markers CD31 and CD144 after 12 days culture of CCPs in Vasculife LS1020 medium and immunolabeling (bottom left panel) for CD31. **F)** Dissociated day 12 cells in Vasculife LS1020 plated for one day in undiluted Matrigel form tubes demonstrated by phase contrast micrograph.

To determine the potency of CCPs to differentiate into iPSC-derived endothelial cells (iPSC-ECs), thawed CCPs were plated in an endothelial medium, Vasculife LS1020^TM^. After 12 days in culture, flow cytometry demonstrated that 88.0±1.3% of cells were CD31^+^/CD144^+^ iPSC-ECs (**Figure 3F**). Further characterization of the CD31^+^/CD144^+^ population shows that 15.2±1.2% of the cells express CXCR4, a marker of arterial endothelial cells (**Figure S1)**.^27,28^ Immunolabeling of the day 12 cultures also demonstrated expression of CD31 in a monolayer of human iPSC-ECs (**Figure 3F)**. To test the functional properties of the iPSC-ECs, a matrigel tube formation assay was carried out on replated day 12 cells. One day after replating, the iPSC-ECs organized into an interconnected network of tubes in Matrigel (**Figure 3G)**. Thus, in medium optimized for EC culture, CCPs readily differentiate to iPSC-ECs.

### CCPs engraft in swine infarcted myocardium and differentiate to cardiac lineages

A total of 70 pigs underwent LAD balloon occlusion reperfusion, of which 52 survived the post-MI period to treatment (**Figure S2**). Of the remaining 52 pigs, 6 pigs were evaluated in the roll-in feasibility cohort which underwent open-chest epicardial injections of CCPs+cECM after 4 weeks post-MI. Pigs were sacrificed 2 weeks later, and hearts were sectioned for histopathology. The other 46 pigs were randomly assigned to one of four treatment groups: CCP+cECM, CCP alone, cECM alone or Control (5% Flexbumin). Entry of pigs into the cECM alone cohort was stopped after only 3 pigs were treated due to lack of a clear trend toward improved ejection fraction and limitations on animal numbers. Thus, the functional studies focused on comparing 3 groups: CCPs, CCPs+cECM, and control.

In the feasibility roll-in cohort, pigs were followed for 2 weeks post cell delivery, and the presence of engrafted human cells was assessed by immunolabeling for human nuclear antigen, Ku80, or human mitochondria. All 6 pigs demonstrated Ku80 positive human cell grafts that were composed largely of cTnI co-labeled cells; however, cTnI labeling intensity was less than observed in the native pig myocardium (**Figure 4A**). Trichrome staining of an adjacent section revealed the corresponding human cell grafts adjacent to the native myocardium (**Figure 4B**). Human cell grafts were identified within the scar and in adjacent native myocardium (**Figure 4C-E**). The grafts were composed predominantly of human cardiomyocytes based on the cTnI immunolabeling, but the sarcomeric pattern and density of labeling was consistent with immature cardiomyocytes. In addition, using co-labeling with an antibody detecting human mitochondria and an antibody for CD31, we detected human endothelial cells in the grafts (**Figure 4F-I**). These results demonstrate that the CCPs can survive following injection into the post-MI porcine heart and differentiate into cardiomyocytes and endothelial cells to form cardiac tissue grafts.

**Figure 4.**
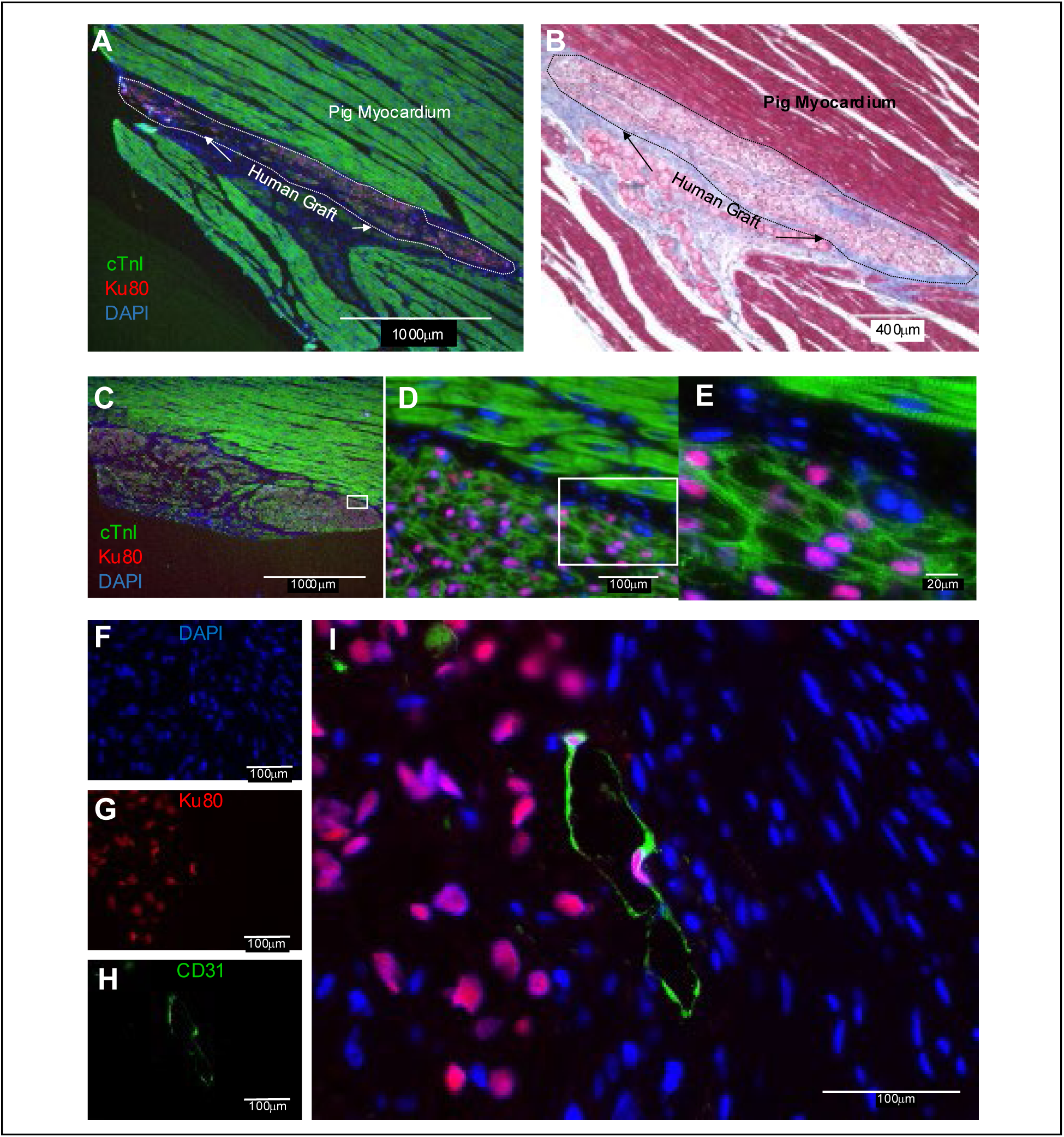
iPSC-CCPs engraft and differentiate to cardiomyocytes and endothelial cells 2 weeks after delivery in post-MI swine heart. **A)** Epifluorescence image of porcine heart section with human graft identified by Ku80 immunolabeling and cardiomyoctyes by cTnI immunolabeling. **B)** Light microscopic image of adjacent histological section stained with Mason’s Trichrome. **C-E)** Immunofluorescence images of human graft in porcine myocardium immunolabeled as indicated identifies human cardiomyocytes. **F-I)** Immunofluorescence image of another human graft in porcine myocardium labeled with anti-Ku80 and anti-CD31 to identify human endothelial cells in the graft.

In the randomized phase, pigs were sacrificed at 4 weeks post-treatment, although a small subset of pigs were survived up to 8 weeks after transendocardial delivery (**Supplemental Table 6** for summary of randomized pigs and analysis). In pigs receiving CCPs with or without cECM, hearts were systematically blocked, and sections were immunolabeled for human Ku80 and cTnI. Human grafts were identified in 13/14 (92.9%) of pigs in each the CCPs and CCPs+cECM cohorts. The graft area observed on the histological sections were comparable between the CCP and CCP+cECM groups (0.68 +/− 0.25 mm^2^ and 0.37 +/− 0.168 mm^2^, respectively, p=0.30, supplemental **Figure S4D**).

Furthermore, the number of graft areas detected was comparable between the CCP and CCP+cECM groups (5.8±2.2 and 5.2±1.6, p=0.57, **Supplemental Figure S4E**) Macroscopic graft size was not measured given the blocking strategy used to enable detailed histological characterization of the large porcine hearts. In the animals sacrificed 4 weeks after CCP delivery, we quantitatively analyzed the cellular composition of the grafts by co-labeling for lineage specific markers and Ku80. The vast majority of Ku80^+^ cells co-labeled with cTnI (∼90%, **Figure 5A,D**) indicating cardiomyocytes. Less than 1% of the Ku80^+^ cells were endothelial cells based on CD31 labeling (**Figure 5B,D)** and 1-2% of the Ku80^+^ cells were vimentin positive stromal cells (**Figure 5C,D)**. Both CCP and CCP+cECM groups showed similar cellular compositions of the grafts. A small fraction (<10%) of the cellular composition of grafts could be other cell types not evaluated for including possible residual CCPs, which were not assessed given the lack of a unique marker to distinguish them from native myocardium. There were no areas of osteochondral differentiation.^29^ Grafts were present in both CCP and CCP+cECM animals up to the latest time point sampled, 8 weeks after delivery.

**Figure 5.**
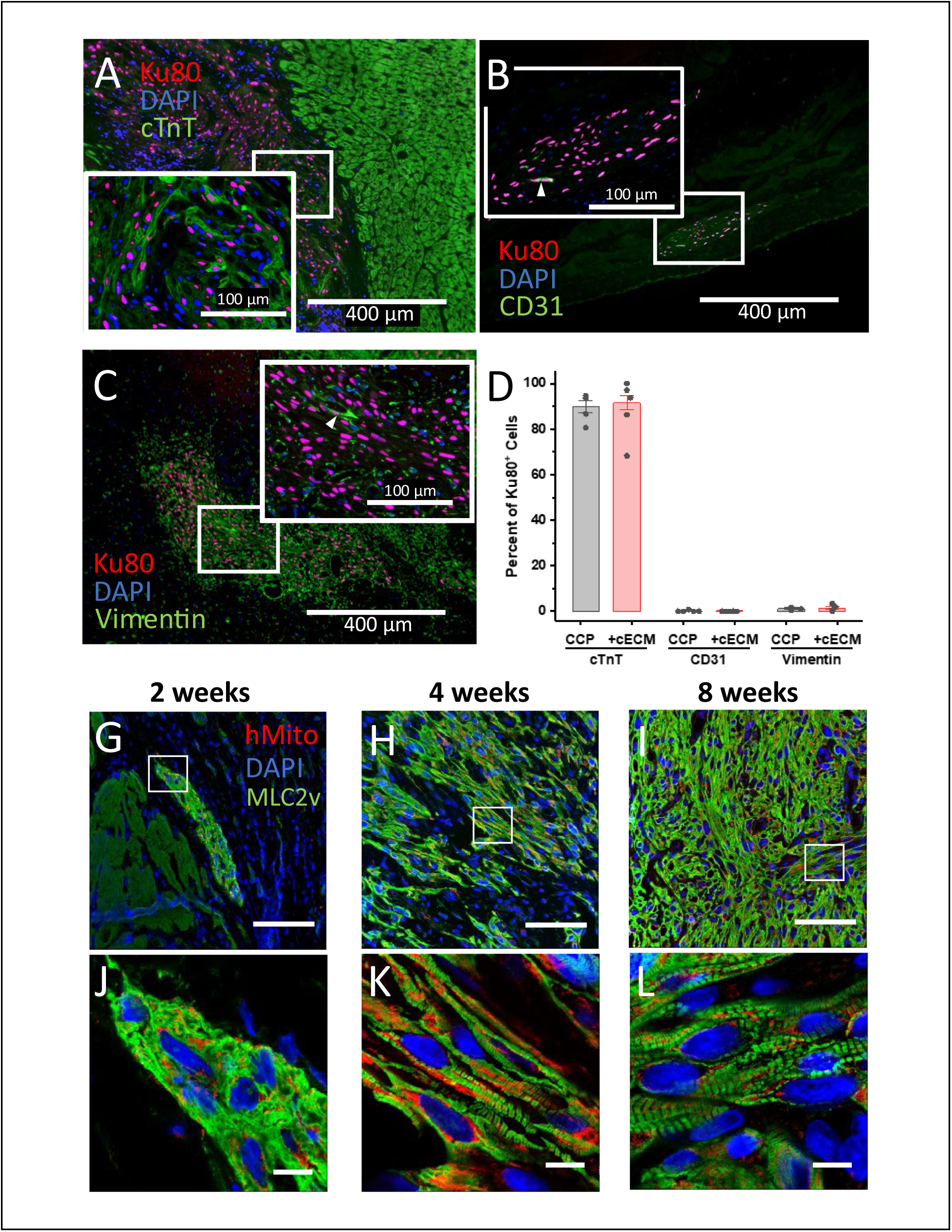
CCP graft composition and maturation. **A)** Immunofluorescence image of 4-week CCP graft dual immunolabeled for cTnT (cardiomyocytes) and Ku80^+^ to detect human cells with DAPI nuclear labeling. **B)** Immunofluorescence image of 4-week CCP graft dual immunolabeled for human CD31 (endothelial cells) and Ku80^+^ with DAPI indicating nuclei. Arrowhead indicates example of human endothelial cells (CD31^+^ and Ku80^+^). **C)** Immunofluorescence image of 4-week CCP graft dual immunolabeled for vimentin (stromal/fibroblasts cells) and Ku80^+^ to identify human cells. Arrowhead indicates example of human stromal cell (vimentin^+^ and Ku80^+^). **D)** Summary data for graft composition of cardiomyocytes (cTnT), endothelial cells (CD31), and stromal/fibroblast (vimentin) per animal (% of total Ku80^+^ cells) at 4-weeks after CCP or CCP+cECM delivery. Bar is average and error bars are SEM, n=4-9. There are no significant differences between CCPs and CCP+cECM for any of the markers. **G-L)** Confocal immunofluorescence images of CCP-derived grafts at time points indicated post-delivery immunolabeled for ventricular-specific MLC2v and human-specific mitochondrial antibody with DAPI to identify nuclei. There is progressive maturation of the MLC2v myofilament pattern with increased organization and density of myofilaments with time. Upper row scale bar 200 µm, and lower row expanded inset from indicated region in white box with scale bar 10 µm. Individual channels for panels J,K,L are in supplemental Figure S5.

### CCP-derived grafts show progressive maturation of ventricular cardiomyocytes

To assess the chamber-specificity of the differentiated cardiomyocytes in the grafts and to examine for evidence of maturation, we immunolabeled sections with an antibody recognizing the ventricular-specific myosin light chain, MLC2v at 2-, 4– and 8-weeks post-transplant. Most of the human cells identified using an anti-human mitochondrial antibody expressed MLC2V at all time points tested (**Figure 5G-L, Figure S5)**. There was evidence of progressive sarcomere organization and maturation based on the pattern of MLC2v labeling from 2 weeks to 8 weeks, although the cells remained smaller and with less sarcomere organization than adjacent adult porcine myocardium. Together these results demonstrate that transendocardial delivery of CCPs in the post-MI swine heart leads to formation of cardiac grafts predominantly composed of human ventricular cardiomyocytes with occasional human endothelial cells and stromal cells. The grafts persist for the 8 weeks of observation in this immunosuppressed swine model and show evidence of progressive cardiomyocyte maturation.

### Cardiac MRI assessment

Twenty-two pigs were evaluable for MRI in the three experimental groups, whereas 19 pigs were excluded from the MRI analysis due to: LVEF >45% (n=1), infarct size <8% of LV (n=1), hemodynamically significant opportunistic sepsis/infection (n=13), and technical issues with the MRI acquisition (n=4) (see **Supplemental Figure 1, Supplemental Table 6**). Prior to CCP delivery, there was no significant difference in baseline post-MI left ventricular ejection fraction: CCPs alone (n=5) 43.8 +/− 4.3%, and CCPs+cECM (n=9) 39.9 +/− 4.1%, and Control (n=8) 44.8 +/− 4.2%. Four weeks following transendocardial delivery, no statistically significant change in ejection fraction was observed for any of the three groups, but CCPs trended to improvement (p=0.08), while CCPs+cECM (p=0.3) and control injections (p=0.99) showed no clear improvement (**Figure 6A,B)**. Similarly, there was no statistically significant difference between groups in left ventricular end-diastolic volume and left ventricular end-systolic volume following intramyocardial injection (**Figure 6C,D)**. Late gadolinium enhancement-determined infarct volume was not significantly changed in any of the three groups (**Figure 6E,F)**.

**Figure 6.**
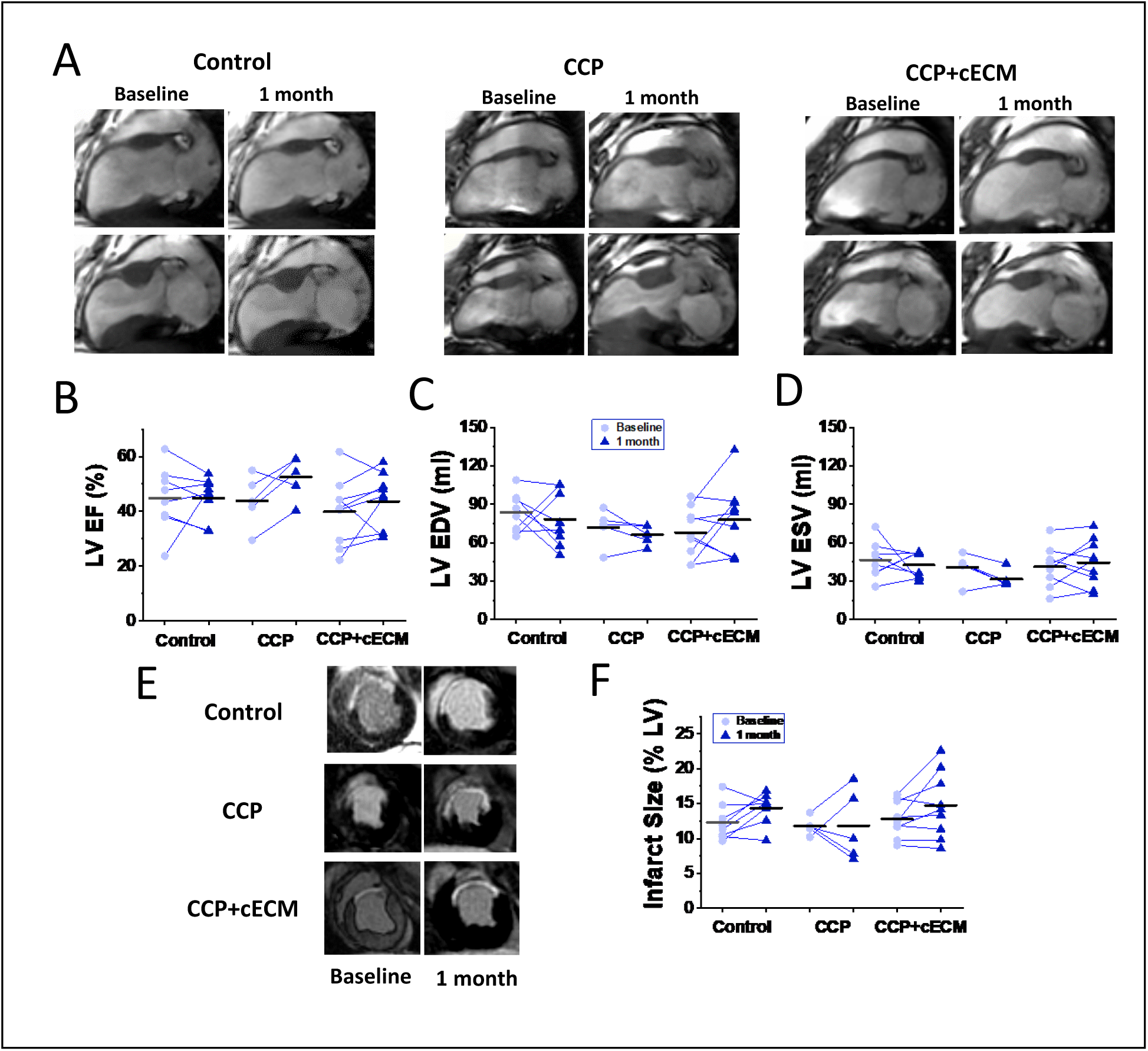
Cardiac MRI analysis. **A)** Representative diastolic (upper) and systolic (lower) cardiac cine MR images from Control, CCP only and CCP+cECM treated animals. **B-D)** Comparisons of baseline and 4 weeks after treatment of LVEF, LVEDV and LVESV showed no statistically significant differences between groups. **E)** Representative cardiac MRI viability images. **F)** Comparisons of baseline and 4 weeks MRI infarct size showed no significant differences. CCP = committed cardiac progenitor cells, cECM = cardiac extracellular matrix, LVEF = left ventricular ejection fraction, LVEDV = left ventricular end diastolic volume, LVESV = left ventricular end systolic volume.

### Pressure-volume hemodynamic analysis shows CCPs improve EF and contractility reserve

Invasive hemodynamic pressure-volume (PV) studies were performed at the endpoint of the study for each animal (**Figure 7A-C)**. Only pigs meeting the MRI inclusion criteria above (EF<45% or infarct >8% of LV, free of infection) and sacrificed at the 4-week timepoint were considered potentially evaluable by PV loop analysis. Technical problems during PV-loop acquisition or poor-quality loops caused the exclusion of 3 additional animals. Comparison of resting ejection fractions from the hemodynamic studies demonstrated a significantly higher ejection fraction in the CCP (46.1 ±1.8%) group compared to Control (37.1 ±2.0%) (p=0.007), and the CCP+cECM (43.3 ±1.3%) group showed a trend to improvement (**Figure 7D**). End systolic volume was significantly reduced in CCP (49.1 +/− 3.0 ml) and CCP+cECM (51.3 +/− 1.7 ml) groups compared to Control (75.3 +/− 5.6 mL) (p=0.0004 ANOVA; p=0.001 and p<0.0001 respectively) (**Figure 7E**). End diastolic volume was also significantly reduced in CCP (91.0 +/− 5.1 ml) and CCP+cECM (90.5 +/− 2.9 ml) groups compared to the Control group (119.3 +/− 6.0 ml) (p=0.0006 ANOVA; p=0.003 and p=0.001 respectively) (**Figure 7F)**. There was no significant difference in end systolic elastance between CCP, CCP+cECM and Control groups at baseline (p=0.13 ANOVA). Following dobutamine challenge to assess contractile reserve, a significant increase in elastance was observed in all groups (CCP alone (8.4 +/−0.9) and CCPs+cECM (4.5 +/− 0.7) control (4.4 +/− 0.7) (compared to baseline (p=0.0002, p=0.004, and 0.04 respectively) (**Figure 7G)**. The CCP alone group showed significantly greater elastance in the presence of dobutamine relative to Control (p<0.006) and CCP+cECM (p=0.007).

**Figure 7.**
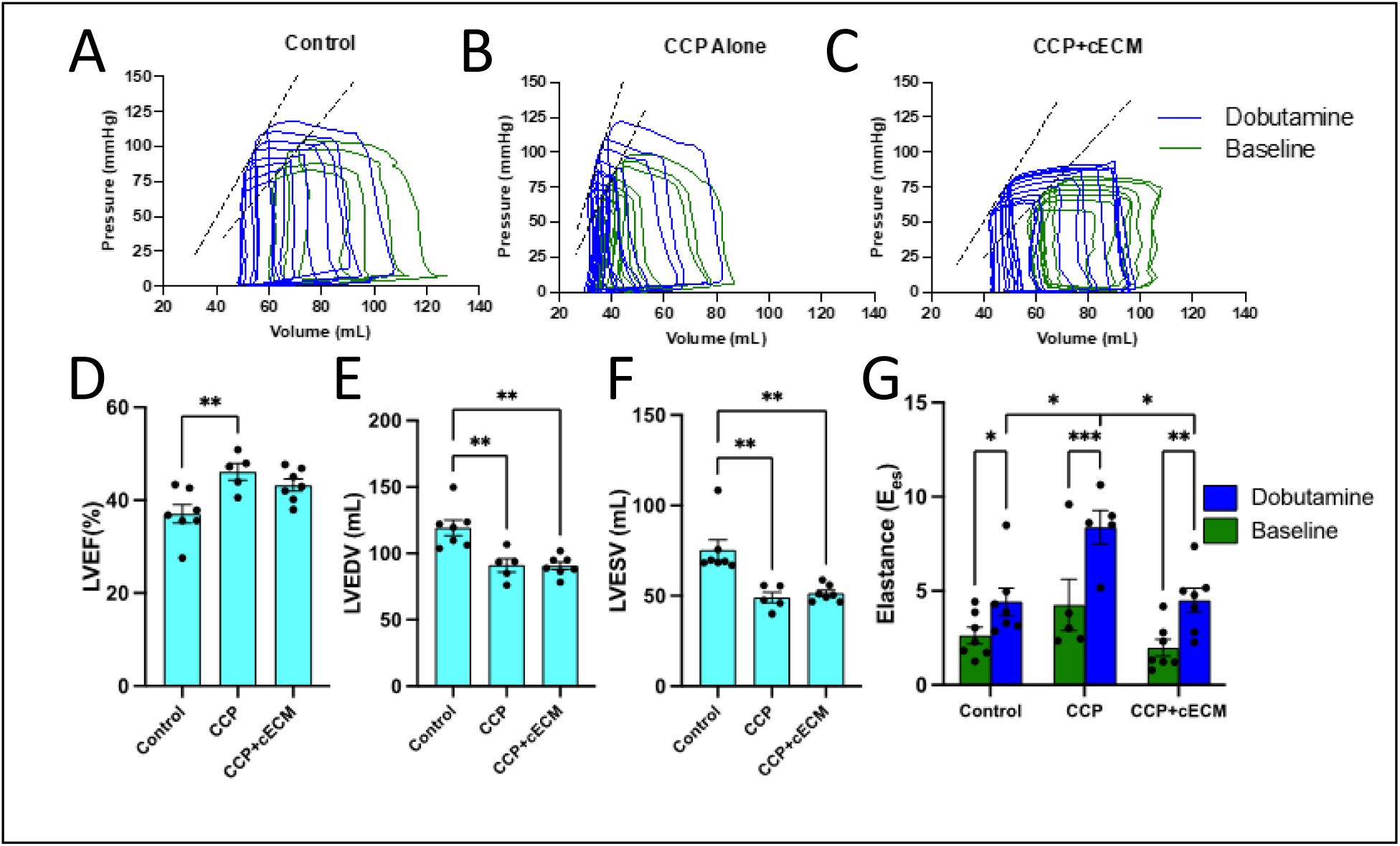
Invasive hemodynamic analysis demonstrates CCP and CCP+cECM show reduced left ventricular volumes and improved contractile reserve. A-C) Representative pressure-volume loop recordings of baseline-dobutamine stress from Control, CCP and CCP-cECM treated pigs at 4 weeks following treatment. **D)** LVEF was significantly greater in the CCP group compared to Control. **E)** LVEDV was significantly reduced in CCP and CCP+ECM relative to Control. **F)** LVESV was significantly reduced in CCP and CCPP+ECM groups relative to Control. **G)** End systolic elastance determined from pressure-volume recordings at baseline and after dobutamine show increased contractile reserve in the CCP and CCP+cECM groups. CCP = committed cardiac progenitor cells, cECM = cardiac extracellular matrix, LVEF = left ventricular ejection fraction, LVEDV = left ventricular end diastolic volume, LVESV = left ventricular end systolic volume *p≤0.05, **p≤0.01, ***p≤0.0001

### Safety evaluation shows no CCP-induced ventricular tachycardia

Safety assessment for the study included evaluation of procedure related complications, gross or histological tumor formation, and arrhythmias. In all treated pigs, there were no procedure related adverse events such as cardiac perforation, MI, stroke, or limb ischemia. Continuous electrocardiographic monitoring for significant tachycardia (>150 bpm for greater than 24 beats) using an implanted cardiac monitor (REVEAL LINQ™) was performed from the time of the initial MI until pigs were sacrificed, in both the initial feasibility cohort and the randomized cohort. There was only one episode of sustained ventricular tachycardia (>30 seconds) that self-terminated in one randomized cohort pig treated with CCP+cECM on post-injection day 2 (**Figure 8B**). This animal (4680) also exhibited post-MI ventricular tachycardia prior to cell delivery. There was one brief run of ventricular tachycardia (10 s) detected in a CCP treated pig on post-injection day 11 (**Figure 8 A,E**). Otherwise, there were no episodes of ventricular tachycardia or ventricular fibrillation detected in any of the pigs treated with CCPs followed for up to 8 weeks. Immunolabeling for the gap junction protein, connexin 43 (Cx43), did show relatively sparse expression of Cx43 within the graft relative to native myocardium and occasionally between graft cardiomyocytes and native myocardium (**Figure 8F)**. Finally, necropsy did not reveal evidence of gross tumors in any animal, and there was no cardiac histological evidence of tumor formation observed.

**Figure 8.**
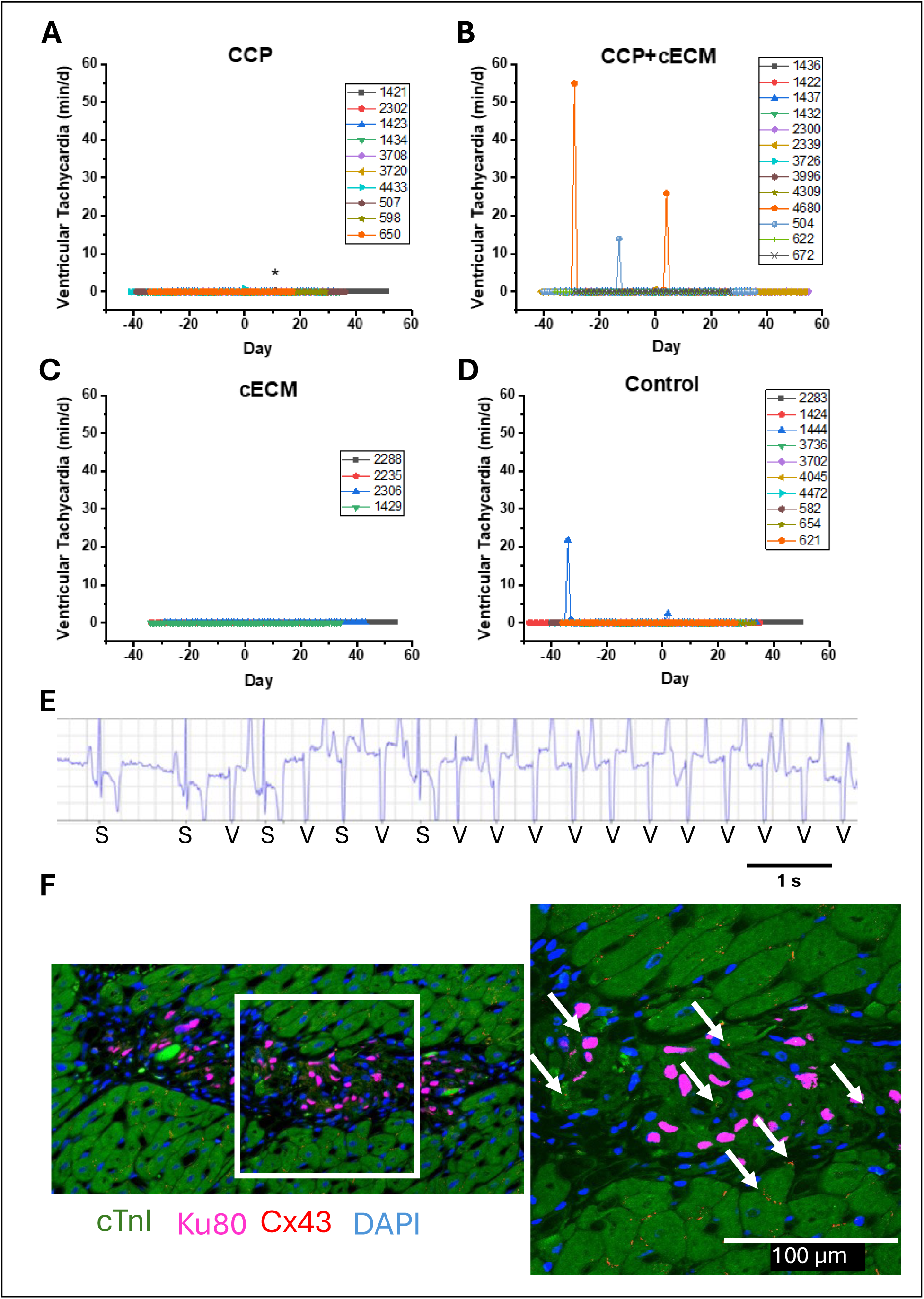
CCP delivery does not result in ventricular tachycardia. **A-D**) Plots show the daily cumulative time each animal is in ventricular tachycardia (defined as >24 beats at rate >150 bpm) per day for each of the 4 randomized groups with recordings starting the day of MI and day 0 being the day of treatment. **E)** Representative ECG recording of the onset of a 10 s run of ventricular tachycardia detected animal 507 on day 11. S indicates sinus complex and V indicates ventricular complex. CCP = committed cardiac progenitor cells, cECM = cardiac extracellular matrix. **F)** Confocal immunofluorescence image of a graft demonstrating Cx43 labeling within host myocardium, in graft and between graft and host using antibodies to against cTNI, Ku80, and Cx43 with DAPI nuclear labeling. Region in white box is at increased magnification to right.

## DISCUSSION

In this preclinical study, we tested the ability of minimally invasive TEI delivery of a novel human iPSC cell product, CCPs, alongside a retention biomaterial to remuscularize the porcine post-MI myocardium. The major findings are: i) transplanted CCPs differentiate to form grafts composed primarily of cTnI and MLC2v positive ventricular cardiomyocyte and occasional human CD31 positive endothelial cells, ii) neither transplant of CCPs or CCPs+cECM significantly improved cardiac MRI measured ejection fraction or cardiac volumes, but the CCP and CCP+cECM groups demonstrated significantly greater contractile reserve relative to control by invasive hemodynamic assessment, iii) CCP +/− cECM administration was feasible without adverse safety outcomes including no tumor formation and no ventricular arrhythmias, and iv) human iPSC-CMs in grafts displayed progressive maturation but remained structurally immature at 8 weeks post injection. Together, these results demonstrate that CCPs hold promise as a therapy for the post-MI heart.

The concept of using cardiac progenitor cells for post-MI heart failure treatment has been investigated in a variety of contexts. The adult heart does not contain a pool of stem or progenitor cells with potency to generate new heart muscle,^30,31^ but hPSCs provide an accessible and clinically relevant source of cardiac progenitor cells. However, hPSC-derived cardiac progenitor cell populations are heterogenous and dynamic during differentiation akin to cardiac development in which different progenitor populations gradually give rise to distinct heart regions and structures. An early study using *in vitro* engineered heart tissue comparing mouse ESC-CMs and ESC-derived Flk1^+^/PDGFRα^+^ cardiac progenitor cells demonstrated that the cardiac progenitor cells differentiated to CMs and improved the electrical coupling in the engineered heart tissue more effectively than the ESC-CMs.^32^ Using similar cell preparations from hESCs, a study comparing hESC-CMs to KDR^+^/PDGFRα^+^ cardiovascular progenitor post-MI in a nude rat model showed comparable grafts and functional effects from the two populations.^33^ A developmentally earlier mesodermal SSEA1^+^ cell population derived from rhesus ESCs was shown to engraft and differentiate to cardiomyocytes, populating 20% of the scar region in the post-MI rhesus heart.^34^ However, a more recent study of the xenogeneic transplantation of hESC-derived SSEA1^+^ cells post-MI nonhuman primate hearts found no human cells remaining at 140 days, although an improvement in cardiac function was observed and attributed to decreased apoptosis of endogenous cardiomyocytes.^35^ Likewise, transplantation of hESC-derived SSEA1^+^ cells in fibrin patches in a rat MI model showed improvement in LV function despite lack of survival of the transplanted cells, suggesting a paracrine mechanism of action.^36^ Studies using an intermediate stage cardiac progenitor population first identified by high levels of Isl1 expression showed in rhesus heart slice models that the cardiac progenitors are highly migratory and can track to areas of cryoinjury.^37^ In post-MI porcine model, the same cardiac progenitors engrafted and formed cardiomyocyte rich grafts, but there was no statistically significant improvement in MRI measures of LV function after 12 weeks.^37^ Yap et al.,^19^ provided evidence that late-stage or committed cardiac progenitors can be effectively delivered at the time of surgically induced MI, engraft and differentiate to relevant cardiac lineages forming cardiomyocytes rich grafts lasting at least 2 months with evidence for improvement in EF from cardiac MRI.^19^ However, the study by Yap et al., found evidence for ventricular arrhythmias following cell delivery in approximately half of the treated animals.^19^ In the present study, we used late stage CCPs, but unlike prior studies they were differentiated from hiPSCs rather than hESCs. Furthermore, although scRNA-seq analysis showed heterogeneity in our CCP preparation, more than 80% of the cells were cardiomyocyte-fated in contrast to the Yap et al. which identified 33% of the cells as cardiac progenitors.^19^ The present study used clinically relevant TEI delivery one month following infarct in contrast to prior studies using surgical, epicardial injection in the acute or subacute phase of MI. Comparable to earlier studies, we observed engraftment and differentiation to form cardiac muscle grafts, but there was not an improvement in cardiac function on cardiac MRI evaluation despite an improvement in dobutamine recruitable functional reserve. The mechanism underlying the improvement in functional reserve could reflect a less advanced stage of heart failure in which β-adrenergic signaling is less desensitized which may be due to a paracrine mechanism or a direct contribution of engrafted cells unmasked by β-adrenergic stimulation. In summary, studies using hPSC-derived cardiac progenitor cells in animal models post-MI demonstrate the ability of the cells to engraft and differentiate with variable evidence for improvement in cardiac function and risk of ventricular arrhythmias.

Significant safety concerns due to ventricular arrhythmias following delivery of hPSC-CMs by epicardial injection in large animal post-MI heart failure models have arisen, although the arrhythmias are generally transient, starting within 1 week after transplant and lasting approximately one month.^11,15,17^ Current evidence suggests that these ventricular arrhythmias are the result of rapid intrinsic automaticity arising from a subpopulation of engrafted CMs rather than reentry mechanisms.^13,16^ The sustained ventricular tachycardia has been tolerated in nonhuman primate models following cell therapy, but in porcine models, there has been significant mortality observed in treated animals.^11,38^ Pharmacological treatment with ivabradine and amiodarone reduced the engraftment arrhythmia burden but did not eliminate it.^38^ Given that abnormal automaticity is the presumed basis for these ventricular arrhythmias, gene editing was used to alter expression of 4 different ion channels and transporters to eliminate automaticity.^14^ This approach greatly reduced engraftment arrhythmias, although there was still an early period of sustained ventricular tachycardia. The arrhythmic risk for cardiac progenitor cells is less clear as some of the large animal studies did not directly monitor for arrhythmias.^34^ However, Yap et al. observed that approximately half of the treated pigs with CCPs developed sustained ventricular tachycardia starting 9 days or later and lasting at least a month and in one animal up to week 12, which appears slightly delayed relative to the hPSC-CM studies. This potentially reflects the additional time needed for differentiation of CCPs. In contrast, the present study observed one relatively short episode of nonsustained ventricular tachycardia in all of the CCP and CCP+cECM treated animals which was in an animal exhibiting runs of ventricular tachycardia prior to cell therapy. The mechanisms underlying the minimal arrhythmic burden in the present CCP study will require further investigation, and potential considerations include differences in the cell population, TEI vs. epicardial delivery, graft size or localization, and time after MI for therapy. Our working model is that gradually differentiating CMs from the CCPs integrate into the native myocardium in smaller, distributed endocardial grafts entrained by the surrounding myocardium without coalescing into large grafts. Future mechanistic studies of electrical integration of the predominantly endocardial cellular grafts resulting from TEI will require alternative approaches from previously successful surface optical mapping of whole heart epicardial grafts.^16^ Furthermore, differences in arrhythmia surveillance approaches may contribute to some differences in observed arrhythmia burden between studies as investigations using implanted loop recorders such as this one, rather than continuous telemetry, will not detect slower ventricular tachycardia (<150 bpm in this study). Ultimately identifying the key factors that eliminate or minimize the arrhythmia risk is necessary for this therapy to advance successfully to the clinic.

Combining cell products with matrix preparations or synthetic hydrogels has been demonstrated in previous studies to improve cell retention and survival in post-MI hearts. For example, the pro-survival cocktail used in several studies includes the complex matrix preparation, Matrigel^TM^.^5^ Gelatin hydrogels have been used successfully in the transplantation of iPSC-CM cardiomyocytes spheroids in post-MI animal models.^39^ In this study, we tested the addition of an injectable formulation of cECM to the CCP product, but this did not result in improved functional benefits at the cECM dose delivered; however, we did find an increase in the number of epicardial grafts on the surface of the heart in the cECM+CCP group relative to CCP alone. This may be related to cECM helping retain any cells that get into pericardial space. How this potential differential localization of grafts impacts long-term outcomes will require additional study. In addition, cECM has been proposed to have favorable immunomodulatory effects via alternatively activated macrophage education,^21^ which could theoretically make local, ischemic tissue more hospitable for co-transplanted cells. However, these favorable influences may be blunted in this immunosuppressed animal model. Overall, the tested delivery of cECM did not improve functional outcomes in this study, but additional studies are needed to determine the optimally formulated and dosed delivery vehicle to promote cell survival and engraftment.

TEI delivery has emerged as an appealing, minimally invasive approach for direct intramyocardial transplantation of cells, biomaterials and other biologic agents in a targeted fashion. In a recent pooled analysis, this delivery approach was observed to be overall safe in >1800 patients injected in clinical trials worldwide^40^. In this study, the NOGA/Myostar (Biosense Webster/J&J) mapping and TEI catheter platform was used to perform targeted injections; however, this platform was recently discontinued from production by the manufacturer. Alternative cardiac mapping approaches to facilitate accurate targeting with TEI, such as co-registered multimodality imaging, have been described but remain investigational.^41^ Similarly, alternative investigational TEI catheters, such as the helical needle-tipped “Helix” catheter (BioCardia, Sunnyvale, CA), are being actively used in active adult cell therapy trials^42,43^. These novel cardiac visualization methods and catheter designs may pave the way to help inject a new generation of therapeutic cells designed to remuscularize the damaged heart.

Overall, the optimal hPSC cell product and delivery strategy to remuscularize the heart remains to be determined. Even a combination of cell products may be preferable such as hiPSC-derived endothelial cells and hiPSC-CMs to promote cell survival and revascularization.^44^ Ground breaking phase 1 clinical trials are testing hPSC-CMs to treat heart failure employing epicardial injections,^45^ transcatheter injections (NCT04982081) and epicardial patches (NCT04945018, NCT04396899) are underway world-wide as well as a completed trial demonstrating the safety of hESC-SSEA1^+^ progenitor cells transplanted at the time of coronary artery bypass grafting surgery.^18^ Despite these developments, important unanswered questions remain including concerns about immunogenicity, repeat dosing possibilities, electromechanical integration, arrhythmogenicity, and whether minimally invasive transcatheter administration will be feasible for wider adoption of this therapy.

Nevertheless, early phase clinical trials are underway testing the safety of delivering hPSC-CMs in patients with heart failure with reduced ejection fraction either by direct intramuscular injection or transplantation of cellularized sheets to the epicardium [clinicaltrials.gov NCT03763136; NCT04945018; NCT03763136; NCT04696328; NCT04945018].

### Limitations

Following cardiac transplantation of the CCPs, we identified the progeny of the CCPs to predominantly include cardiomyocytes and more rare endothelial cells and fibroblasts; however, we cannot exclude other small populations of cell types including residual CCPs. We did not observe tumor formation which suggests that the cell preparation did not include pluripotent stem cells capable of forming teratomas. The dosing of CCPs was based on the literature and pilot studies of epicardial injected CCPs, so improvements in dosing and repeat dosing could positively impact outcomes. The current study did not provide long-term follow-up of the animals, in part due to the challenge of opportunistic infections in the setting aggressive immunosuppression. There was qualitative evidence of maturation over 2 months based on myofilament organization, but no quantitative measure of maturation was made. Infections also caused a reduction in the planned sample size. Electrical integration of the cells was not directly examined in this study, but it will be an important subject of future research to define the mechanisms leading to lack of arrhythmias in our study relative to hPSC-CMs studies.

## Acknowledgements

Sincere thank you to Allison Rodgers, Everett Temme, Sarah El Meanawy and Vivian Hacker for animal handling and procedures, and Emily Ferris for MRI acquisition and analyses.

