## Supplement 1 for "Human iPSC-derived Committed Cardiac Progenitors Generate Cardiac Tissue Grafts in a Swine Ischemic Cardiomyopathy Model without Triggering Ventricular Arrhythmias"

**SUPPLEMENTAL METHODS**

<u>Committed Cardiac Progenitor Cells</u>

Cryopreserved human iPSC-derived CCPs were obtained from Fujifilm Cellular Dynamics, Inc. (FCDI), Madison, WI. Briefly, the therapeutic grade iPSC line 31537.101 was reprogrammed from peripheral blood mononuclear cells (PBMCs) using proprietary FCDI technology and expanded to large master and working cell banks. Differentiation of iPSCs to CCPs was achieved by modulating the Wnt/beta-catenin and TGFbeta/Actinin pathways using small molecules (CHIR99021, XAV939, SB431542) at specific timepoints during the process that included aggregate formation of iPSCs, mesoderm induction, and specification towards a cardiac progenitor cell-fate (See Figure 1A). In brief, working cell banks were thawed into and expanded in multiple vitronectin-coated (ThermoFisher Scientific) CellSTACK® cell culture chambers (Corning), harvested with TrypLE (ThermoFisher Scientific), resuspended in Essential 8 Medium (ThermoFisher Scientific) containing 1.0 μM H1152 (EMD Scientific) and 2.0 μM CHIR99021 (Biovision) and $1X10^6$ cells/ml were seeded into 3L PBS3 MagDrive Bioreactors (PBS Biotech, Inc.) to initiate differentiation for large-scale cardiac progenitor cell manufacturing, stirring at 25 rpm. After 1 day, the aggregates were allowed to settle before replacing 80% of the medium with RPMI (ThermoFisher Scientific) plus B27 supplement without insulin (1x) (ThermoFisher Scientific) and 4.4 μM CHIR99021. On day 3, 80% of the medium was changed to RPMI plus B27 supplement with 10 μM XAV939 and 2 μM SB431542. On day 4, the medium was changed to RPMI plus B27 supplement. On day 6, approximately 4-6 x $10^9$ CCPs were harvested from each bioreactor, dissociated with TrypLE, and cryopreserved with CryoStor CS10 (Biolife Solutions) at $300x10^6$ cells/5mL in AT-Closed

Vials (Aseptic Technologies) using a controlled rate freezer and stored long-term in the vapor-phase of liquid nitrogen tanks.

### *In vitro* differentiation potential of CCPs

CCPs were thawed directly into RPMI (ThermoFisher Scientific) + B27 supplement (1x) (ThermoFisher Scientific), washed, and resuspended in either RPMI+B27 for cardiomyocyte differentiation or VascuLife LS1020 (LifeLine Cell Technology) for endothelial differentiation. For cardiomyocyte differentiation, CCPs were plated into vitronectin-coated 6-well plates at $3x10^6$ cells/2mL/well or vitronectin-coated 96-well plates at $5x10^4$ cells/200uL/well. For endothelial differentiation, CCPs were either plated into diluted Matrigel coated T225 flasks at $3.4x10^6$ cells/45mL/flask or diluted Matrigel coated 12-well plates at $5x10^5$ cells/1mL/well. Respective cardiac or endothelial medium was refreshed 24hr post plating and every other day there-after until day 7 where the cells were washed with DPBS(-/-), and either fixed for immunocytochemistry or harvested with TrypLE to be re-plated or analyzed by flow cytometry.

### Tube Formation assay

Day 7 CCP-derived endothelial cells were plated into undiluted Matrigel coated 24-well plates at various densities in 400 µl of Vasculife media/well. After 24 hr, cultures plated at 100k/well were imaged for tube formation using Nikon Eclipse Ts2 microscope with 4x objective.

### Flow Cytometry

Cells were immunolabeled directly after thaw or after being harvested from *in vitro* differentiation assays. Cell surface protein expression was performed by labeling $2x10^5$

cells for KDR-PE (R&D Systems, FAB357P) + PDGFRα-APC (Alexa Fluor 647?) (Cell Signaling, 5876S); CD56-APC (BD Pharmingen, 555518) + CXCR4-PE (BioLegend, 306506); EpCAM-FITC (Miltenyi, 130-113-263); CD31-FITC (BD Pharmingen, 555445) + CD144-647 (BD Pharmingen, 561567); CD140b-PE (BD Pharmingen, 558821) + CD90-APC (BD Pharmingen, 559869) diluted in wash buffer (DPBS(-/-)+2% FBS) at 4°C for up to 60 minutes followed by washing and analysis with a BD Accuri flow cytometer. Intracellular protein expression was performed by staining $2x10^6$ cells with Live/Dead Green stain (Molecular Probes, L23101) for 15 minutes, washed, then fixed for 15 minutes with 4% formaldehyde. Fixed cells were washed twice with FACS permeabilization buffer (DPBS(-/-) + 0.1% saponin) and labeled with mouse anti-sarcomeric alpha-actinin (Abcam, ab9465) for at least 1 hour at 4°C. Cells are then washed twice with FACS permeabilization buffer, immunolabeled for 30 minutes with Alexa Fluor 647 donkey anti-mouse IgG (H+L) (ThermoFisher, A-31571), washed twice and analyzed with a BD Accuri flow cytometer.

Immunocytochemistry

Immunocytochemistry analysis for differentiation of CCPs to cardiomyocytes *in vitro* was performed using cells cultured in 96-well plates. Briefly, cells were washed with DPBS(-/-), fixed with 4% formaldehyde for 15 minutes and washed with ICC permeabilization buffer [DPBS (-/-) + 0.1% triton-X]. Cells were stained overnight at 4°C with mouse anti-sarcomeric alpha-actinin (Abcam) + rabbit anti-NKX2.5 (Cell Signaling) or mouse anti-cardiac troponin T (ThermoFisher Scientific) + rabbit anti-cardiac troponin I (Abcam). The next day the cells were washed with ICC permeabilization buffer, stained with the appropriate Alexa Fluor donkey anti-mouse and anti-rabbit secondary antibodies for 30 minutes followed by nuclei staining with DAPI and imaged with an EVOS cell imaging system (ThermoFisher Scientific).

Immunocytochemistry analysis for differentiation of CCPs to endothelial cells was performed on cells cultured in 12 well plates. Briefly, the cells were washed with DPBS(-/-), fixed with 4% formaldehyde for 15 minutes and washed with ICC donkey perm-block buffer (DPBS(-/-) + (donkey serum) + 0.2% saponin). Cells were then immunolabeled for 4hr at 4°C with sheep anti-CD31 (R&D Systems, AF806) + rabbit anti-cTNT (Abcam, ab45932). After primary incubation, cells were washed with ICC donkey perm/block buffer, labeled with the appropriate Alexa Fluor donkey anti-sheep 647 (Invitrogen, A21448) and anti-rabbit 488 (Invitrogen, A21206) secondary antibodies for 30 minutes followed by nuclei staining with NucBlue (Invitrogen, R37606) and imaged with Molecular Devices High content cell imaging system.

Gene Expression by qPCR

Bulk RNA gene expression was performed in a two-step RT-PCR reaction followed by qPCR using the Fluidigm BioMark HD system. All steps were executed as described in Fluidigm's Real Time PCR Users Guide (68000088), appendix C. Briefly, 200 ng of total RNA prepared above was used as starting material for cDNA first strand synthesis using High Capacity RNA to cDNA kit (Applied Biosystems). The preamplification was done with TaqMan PreAmp Master Mix (Thermo Fisher Scientific) and 48 pooled primer pairs (50nM final concentration each primer) for 10 cycles (15 seconds at 95°C, 4 min at 60°C). Unincorporated primers were removed by Exonuclease digestion for 30 min at 37°C followed 15min inactivation at 80C (New England BioLabs). Samples were diluted 5-fold with DNA suspension buffer (Teknova). The qPCR was performed with Fluidigm's

48.48 dynamic array IFC on the Biomark HD using 2X SsoFast EvaGreen Supermix with Low Rox (Bio-Rad Laboratories), 20X DNA Binding Dye Sample Loading Reagent (Fluidigm), and primers at a final concentration of 500nM. The cycling parameters were as follows: hot start 95°C for 60 seconds, then 30 cycles of 96C for 5 sec, 60°C for 20 sec, followed by 60° – 95°C melting curve. cT values for gene expression values were generated and exported from Fluidigm Real-Time PCR analysis software. Relative gene expression values presented were normalized by RPL32 expression.

### Transcriptomic analysis of CCPs by single-cell RNA sequencing

Cryopreserved CCPs stored in liquid nitrogen were rapidly thawed in a 37 °C water bath for 2 min. The contents of each vial were immediately transferred to a 15 mL conical tube containing 1 mL of thawing medium (PBS with 1% BSA, RT). The empty vial was rinsed with an additional 1 mL of thawing medium and the rinse was combined with the cell suspension to maximize recovery. Cells were washed twice with thawing medium and centrifuged at 300 × g for 8 minutes each. After centrifugation, cells were filtered using a Flowmi™ Cell Strainer (40 µm pore size) to remove aggregates and debris. Cell concentration and viability were measured using a NucleoCounter® NC-200. Samples with ≥70% viability were selected for downstream processing using the 10X Genomics Chromium Next GEM Single Cell 3', targeting recovery of ~10,000 cells per sample. A minimum concentration of 400 cells/µL was required to proceed. Final libraries were pooled according to the 10X Genomics Chromium Next GEM Single Cell 3’ protocol and sequenced on an Illumina NovaSeq 6000 instrument using a kit able to generate ~40,000–50,000 reads per cell.

Chromium single cell data output from 10X Genomics analysis pipeline were further analyzed using Seurat and Cellxgene VIP software (BioInfoRx, Inc., Madison WI). Differentially expressed genes (DEGs) and UMAP were generated from Seurat analysis using the standard resolution. Dot Plot and embedding Plot are generated using the Cellxgnene VIP software. GO term analysis was performed using the open source g:Profiler (https://biit.cs.ut.ee/gprofiler/gost). The input to g:Profiler is the DEGs for each cluster. The output includes enriched terms in MF (Molecular Function), BP (Biological Process), CC (Cellular Component), KEGG and REAC pathway.

Cardiac fibroblast-derived Extracellular Matrix

Briefly, cECM (Tandem™, Cellular Logistics Inc, Madison, WI) was produced from primary human cardiac fibroblasts cultured under high-density conditions followed by a decellularization procedure as previously described.[20,21] The injectable cECM formulation was manufactured by milling cECM scaffolds into a fine particulate of approximately 80-micron diameter using a pharmaceutical bead mill. cECM particles were added to the cell suspension and the cells are allowed to bind at room temperature for 1 hour prior to injection.

Closed Chest Porcine MI Model

All procedures were performed in accordance with a University of Wisconsin-Madison Institutional Animal Care and Use Committee approved protocol, M005512-R01-A05. Closed chest, 120 minute balloon-occlusion and reperfusion of the mid-left anterior descending artery was used to induce large anteroseptal MIs in male and female Yucatan swine (6-7.5 months age, ~30kg, Premier Biosource, MI) using percutaneous methods previously described by our group[24,25]. Amiodarone 25mg/kg po was administered daily starting 5 days pre-MI through 7 days post-MI. Heparin 70 U/kg IV was administered hourly during MI creation in all pigs. Lidocaine 50-100 mg/hr was administered as a constant rate infusion starting just prior to ischemia through to animal recovery. Amiodarone 50-100 mg IV, epinephrine 0.1-0.2 mg IV and external defibrillation at 200 J were administered as necessary to treat acute ventricular arrhythmias. Laboratory personnel and veterinary staff continuously monitored animals for their welfare before, during and after procedures. Pigs were survived 4 weeks post MI at which time they received the investigational treatment.

Experimental Design

There were two phases in the overall study. First, a roll-in, non-randomized cohort of pigs underwent direct epicardial needle injections of CCPs plus cECM into the infarct and peri-infarct region via an open chest thoracotomy to assess feasibility, establish best practices, and optimize immunohistochemistry to localize human cell grafts prior to launching the larger randomized-controlled trial. Following treatment, the roll-in cohort of pigs were survived for 2 weeks and then sacrificed. Arrhythmia surveillance and histopathology was performed; however, MRI and pressure-volume assessment were not performed since the primary objective was feasibility. Thereafter, an additional larger cohort of pigs were randomized by computer-generated block randomization into four treatment groups: 1) CCPs plus cECM, 2) CCPs alone, 3) cECM alone and 4) Control (Flexbumin), where the investigational products were administered by transendocardial catheter injections. Baseline MRI assessment was performed at 4 weeks post-MI followed by treatment. Four weeks after treatment, most pigs underwent cardiac MRI assessment followed by invasive pressure volume hemodynamic evaluation and sacrifice. A small subgroup of pigs were survived to 8 weeks post treatment (**Figure S2 for allocation of pigs)**.

Epicardial Injections (Feasibility Cohort)

Due to the multi-faceted nature of this translational study, a small cohort of immunosuppressed post MI pigs underwent 10 epicardial injections of CCP+cECM into the infarct and peri-infarct border via an antero-lateral thoracotomy. Animals were recovered for 2 weeks, and then sacrificed. Detailed histopathology and continuous ECG recording was performed using an implantable ECG recorder (Reveal LinQ, Medtronic, Minneapolis, MN) that was surgically implanted in the skin adjacent to the left scapula. No MRI or pressure volume loop evaluation was performed in this small, single-arm feasibility cohort as its main purpose was simply to inform manufacturing feasibility, deliverability and to optimize tissue sectioning and immunohistochemistry staining methods.

Transendocardial Injections (Randomized Cohort)

An electroanatomical map of the left ventricle was performed with the NOGA XP Cardiac Navigation System (Biosense Webster) and NOGAstar mapping catheter to create an endocardial roadmap and distinguish viable from non-viable myocardium (**Figure S3, Supplemental Video 1**).[26-28] An 8F electroanatomically tracked, deflectable TEI catheter with a 27G straight hollow needle design (Myostar, Biologic Delivery Systems, Irvine, CA) was used by an experienced operator who was blinded to the treatment assignment to perform the injections (**Figure S3, Supplemental Video 1)**. Prior to initiating the injections, the needle injection depth was pre-calibrated to 3.5 mm when the catheter was placed in a curved in silico aortic arch with a 90-degree deflection. The injection strategy consisted of a total of 14 injections (200 microliter per injection) of study agent with approximately 50% of injections delivered into the peri-infarct borderzone and 50% directly into the infarct itself. Each injection was performed over 30 seconds with a 15-second dwell time.

Dose

Using aseptic technique, CCPs obtained from Fujifilm CDI, Madison, WI were rapidly thawed in a 37°C water bath (ThawSTAR, Biolife Solutions, Bothell, WA). Post-thaw viability, assessed using trypan blue exclusion, was 92.9±2.1% (n=28, mean±S.D.). 5% Flexbumin, which was used to suspend the cells and cECM, was used as a treatment control. Study agents were loaded into three, partially opaque 1 mL Luer lock syringes at room temperature. Intermittent gentle agitation was applied until injection. The total per pig CCP dose was 300 million CCPs and the total cECM dose was 50 mg. Following injection, residual cells were recovered, and measured viability by trypan blue exclusion was 82.6±5.0% (n=14, mean±S.D.) for CCP group and 82.1±5.6% (n=14, mean±S.D.) for CCP+cECM group.

Immunosuppression

To prevent immune rejection of the transplanted human cells, a 4-drug immunosuppression regimen was initiated prior to investigational treatment: i) cyclosporine 10 mg/kg po twice daily with adjustments to achieve trough levels above 250 microgram/L, ii) methylprednisolone 2.5 mg/kg po twice daily, iii) mycophenolate mofetil 250 mg oral twice daily and iv) abatacept 500 mg IV biweekly were administered. Sulfamethoxazole/trimethoprim 960 mg oral twice daily was administered to prevent infections. Blood samples were drawn every 2 weeks following investigational treatment for complete blood count, chemistry panel and cyclosporine levels.

### Arrhythmia Surveillance

Continuous multi-lead telemetry monitoring was performed during MI model creation, investigational treatments, MRI and invasive P-V loop evaluation. An implantable ECG recorder (Reveal LinQ™, Medtronic, Minneapolis, MN) was surgically implanted subcutaneously adjacent to the left scapula at the time of the MI intervention using methods previously described by our group.[24] We previously confirmed that implanting these devices in this location was not associated with MR image distortion, and the magnetic field and radiofrequency did not interfere with ECG recordings.[24] The ECG recorder was programmed to detect: tachycardia>150 bpm for >24 beats, asystole for >4.5 sec, bradycardia <30 bpm for at least 4 beats and atrial fibrillation. Sustained ventricular tachycardia was defined as a run of ventricular tachycardia lasting ≥30 sec. Wireless device interrogation was performed on day 2, and then weekly post-MI. Rhythm strips from every recorded event were interpreted by an experienced clinical cardiologist (TJK) who was blinded to the treatment group assignment.

### Magnetic Resonance Imaging (MRI)

Cardiac MRI was performed prior to treatment at baseline (4 weeks post-MI) and at 4 weeks post-treatment on a 3T Premier scanner (GE Healthcare, Waukesha, WI). Specifically, left ventricular chamber size and function was measured using short axis oblique slices with balanced steady-state free procession (bSSFP) imaging. (Scan parameters: TR/TE/Flip angle=3.3 ms/1.0 ms/50; field-of-view (FOV)/matrix=35×35 cm/200×224; slice thickness/spacing=10 mm/0 mm; cardiac phases/views per segment=20/16). Infarct size was measured using standard delayed enhancement (DE) inversion recovery based sequence performed 10 min following intravenous administration of 0.10 mmol/kg Gd-BOPTA (Multihance, Bracco Diagnostics, Princeton, NJ, scan parameters: TR/TE/TI/Flip angle=4.7 ms/1.0 ms/200–280 ms/25°; FOV/Matrix=35× 31.5 cm/224×192; slice thickness/spacing= 10 mm/0 mm; views per heartbeat=18). The trigger delay was adjusted to image mid-diastole based on the multiphase bSSFP series. All MRI imaging was performed at end expiration by temporarily suspending mechanical ventilation.

Anatomic and functional cardiac image analysis was performed by blinded personnel at the Radius Medical Imaging Analysis Core Laboratory using cvi42, Circle Cardiovascular Imaging, Calgary, CA. Cine bSSFP images were segmented to measure indexed LV end-diastolic volume, end-systolic volume, ejection fraction, stroke volume and radial, longitudinal and circumferential cardiac strain. Infarct size was quantified using late gadolinium enhancement imaging. Inclusion criteria for MRI evaluation was 4 week post-MI ejection fraction of <45% or infarct size of >8% of the LV size and freedom from sepsis or other severe infection.

Invasive Pressure Volume Assessment

Prior to sacrifice, invasive pressure-volume hemodynamic measurements were performed using a 5F high fidelity multi-ring electrode conductance catheter (Scisense Inc, London, ON) positioned retrograde into the left ventricle via transcarotid artery access. A 24 - 28 mm diameter balloon catheter was positioned via femoral vein into the inferior vena cava. Contralateral femoral venous access was obtained to advance a 7F Swan Ganz catheter under fluoroscopic guidance into the pulmonary artery. Calibrating stroke volume estimates were obtained by Swan Ganz thermal dilution method. Pressure–volume data was recorded at steady state and during transient inferior vena cava transcatheter balloon-occlusion, both at pre-dobutamine and with 20 mcg/kg IV dobutamine infusion using previously described methods.[24] Pre-load independent contractility reserve was defined as the difference in end-systolic pressure-volume relationship (ESPVR) between dobutamine stress and rest.

Myocardial contractility and work were indexed by the maximal rate of isovolumetric contraction (+dP/dt), stroke work (SW), and ventricular elastance, slope of the end-systolic pressure–volume relationship (Ees). Pre-load was analyzed as end-diastolic volume and pressure and afterload evaluated as effective arterial elastance (the ratio of LV end-systolic pressure to stroke volume). Diastolic function was indexed by LV end-diastolic pressure and the time constant of ventricular relaxation (Weiss formula). Hemodynamic pressure–volume data was digitized at 200 Hz and stored for subsequent analysis on a personal computer by using commercial software (iWorx Dover, New Hampshire, USA).

Histopathology Assessment

Hearts were excised and hung on a modified Langendorf apparatus and perfused with 1 L heparinized saline followed by 1 L 10% neutral buffered saline. The infarct-related scar was identified grossly and a region largely encompassing the scar from just above the infarct region to the apex was dissected (**Figure S4,** Region A). 8-12 blocks, 10-15 mm each, were cut on the short axis and placed into tissue cassettes. Next, regions on either side of the infarct were dissected (**Figure S4,** Regions B and C) and blocked as described above. In total, approximately 30 tissue blocks, encompassing approximately

75% of the left ventricle were taken per heart. The blocks were placed in 10 % neutral buffered formalin for 24 h before transfer to 70 % ethanol until histological processing and paraffin embedding. All blocks from an individual animal (~30) were then microtome sectioned (5 μm) to produce a representative section from the basal side of each block. Following antigen retrieval (citrate buffer, pH 6.0, 10 mM Citric Acid, 0.05% tween 20), for 3 minutes in a Biocare decloaker (Biocare Medical, Concord, CA), sections were co-immunolabeled for human species selective markers Ku80 and anti-TnI. Sections were analyzed independently by 3 experienced reviewers to identify, localize, and measure the area of human grafts (Ku80$^{+}$) using an EVOS (Life Technologies) microscope. Grafts were categorized as in the infarct zone if they were at least partially surrounded by infarcted related scar or as borderzone if they were completely surrounded by normal appearing myocardium. If no grafts were identified on any of the sections from a given animal, all blocks underwent additional sectioning every 500 μm until human grafts were identified or until the tissue in the block was exhausted. The area of human cardiomyocyte grafts was measured using NIH ImageJ for the first section per block where grafts were identified.

To identify endothelial cells derived from CCPs, fixed tissue sections were co-immunolabeled using antibodies against human CD31 (Zeta Corp., JC70A) and human Ku80 (nuclear marker; Cell Signaling Technology, 2180), followed by an Alexa Fluor 568-conjugated goat anti-mouse (Thermo Fisher Scientific, A-11004) and Alexa Fluor 647-conjugated goat anti-rabbit (Thermo Fisher Scientific, A-21070) secondary antibody.

To identify cells of mesenchymal lineage—including fibroblasts, smooth muscle cells, pericytes, and endothelial cells—derived from CCPs, fixed tissue sections were co-immunolabeled with antibodies against Vimentin (Thermo Fisher Scientific, MA5-11883) and human KU80 (Cell Signaling Technology, #2180), followed by an Alexa Fluor 568-conjugated goat anti-mouse (Thermo Fisher Scientific, A-11004) and Alexa Fluor 647-conjugated goat anti-rabbit (Thermo Fisher Scientific, A-21070) secondary antibody. Vimentin was used as a marker for mesenchymal cells, while KU80 was used to identify human cell origin.

To assess the level of integration between native cardiomyocytes and engrafted cells differentiated from CCPs, tissue sections were immunostained for Connexin 43 (Biorbyt, orb11602), cardiac Troponin I (Abcam, ab10231), and human KU80 (Cell Signaling Technology, #2180), followed by an Alexa Fluor 593-conjugated donkey anti-goat (Thermo Fisher Scientific, A-11058), Alexa Fluor 488-conjugated goat anti-mouse (Thermo Fisher Scientific, A-11001) and Alexa Fluor 647-conjugated goat anti-rabbit (Thermo Fisher Scientific, A-21070) secondary antibody. This combination allowed for the evaluation of cell–cell coupling and identification of engrafted human-derived cells within the myocardium.

**Supplemental Table 1. Flow Cytometry of Differentiation Time Course: EPCAM$^{+}$ %** (Data for Figure 1B)

| *Day* | 1 | 2 | 3 | 4 | 5 | 6 |
|---|---|---|---|---|---|---|
| *MB208 V1 31537_104918* | 99 | 99 | 91 | 57 | 17 | 5 |
| *MB208 V2 31537_105013* | 99 | 99 | 82 | 29 | 5 | 1 |
| *AAK11 1* | | | | 24 | 7 | 3 |
| *AAK11 2* | | | | 26 | 8 | 2 |
| *AAK11 3* | | | | 27 | 8 | 2 |
| *MB204 1* | | | | 30 | 11 | 3 |
| *MB204 2* | | | | 22 | 7 | 3 |
| *MB204 3* | | | | 27 | 9 | 3 |
| *MB204 4* | | | | 33 | 9 | 3 |
| *MB204 5* | | | | 33 | 13 | 3 |
| *MB204 6* | | | | 30 | 8 | 3 |
| *EH94* | | | | | | 12 |
| *EH95* | | | | 46 | 28 | 6 |
| *EH108 1* | 98 | 98 | 87 | 83 | 42 | 7 |
| *EH108 2* | 98 | 98 | 85 | 69 | 33 | 7 |
| *EH111 1* | 99 | 99 | 64 | 37 | 10 | 3 |
| *EH111 2* | 99 | 99 | 84 | 40 | 12 | 2 |
| *EH111 3* | 99 | 99 | 89 | 68 | 23 | 9 |
| *MB235 A* | | | | | | |
| *MB235 B* | | | | | | |
| ***AVERAGE*** | **99** | **99** | **83** | **40** | **15** | **4** |
| ***STDEV*** | **0** | **0** | **9** | **18** | **11** | **3** |
| ***N*** | **7** | **7** | **7** | **17** | **17** | **18** |

**Supplemental Table 2. Flow Cytometry of Differentiation Time Course: CD56$^+$CXCR4$^-$ %** (Data for Figure 1B)

| *Day* | *1* | *2* | *3* | *4* | *5* | *6* |
|---|---|---|---|---|---|---|
| *MB208 V1 31537_104918* | 4 | 6.2 | 5.4 | 30 | 80 | 93 |
| *MB208 V2 31537_105013* | 3 | 5.8 | 7.5 | 59 | 93 | 97 |
| *AAK11 1* | | | 3.6 | 52 | 92 | 94 |
| *AAK11 2* | | | 3.4 | 46 | 92 | 95 |
| *AAK11 3* | | | 3.5 | 50 | 90 | 95 |
| *MB204 1* | | | 11 | | | 99 |
| *MB204 2* | | | 14 | | | 99 |
| *MB204 3* | | | 6 | | | 99 |
| *MB204 4* | | | 12 | | | 99 |
| *MB204 5* | | | 10 | | | 99 |
| *MB204 6* | | | 15 | | | 99 |
| *EH94* | | | 13 | 22 | 55 | 92 |
| *EH95* | | | 9 | 37 | 65 | 84 |
| *EH108 1* | 0 | 0 | 5 | 42 | 63 | 90 |
| *EH108 2* | 0 | 0 | 8 | 35 | 70 | 90 |
| *EH111 1* | 0 | 0 | 30 | 81 | 93 | 98 |
| *EH111 2* | 0 | 0 | 48 | 82 | 92 | 99 |
| *EH111 3* | 0 | 0 | 22 | 58 | 79 | 98 |
| *MB235 A* | | | 21 | 75 | 94 | 98 |
| *MB235 B* | | | 23 | 81 | 94 | 96 |
| ***AVERAGE*** | **1** | **2** | **14** | **54** | **82** | **96** |
| ***STDEV*** | **2** | **3** | **11** | **20** | **14** | **4** |
| ***N*** | **7** | **7** | **20** | **14** | **14** | **20** |

**Supplemental Table 3. Flow Cytometry of Differentiation Time Course: KDR$^{+}$PDGRα$^{+}$%** (Data for Figure 1B)

| *Day* | *1* | *2* | *3* | *4* | *5* | *6* |
|---|---|---|---|---|---|---|
| *MB208 V1 31537_104918* | 0 | 18 | 22 | 42 | 44 | 48 |
| *MB208 V2 31537_105013* | 0 | 18 | 32 | 67 | 27 | 27 |
| *AAK11 1* | | | | 32 | 4 | 2 |
| *AAK11 2* | | | | 54 | 11 | 1 |
| *AAK11 3* | | | | 46 | 8 | 2 |
| *MB204 1* | | | | 64 | 41 | 38 |
| *MB204 2* | | | | 78 | 38 | 38 |
| *MB204 3* | | | | 71 | 36 | 38 |
| *MB204 4* | | | | 70 | 40 | 38 |
| *MB204 5* | | | | 63 | 41 | 38 |
| *MB204 6* | | | | 67 | 36 | 38 |
| *EH94* | | | | 48 | 63 | 4 |
| *EH95* | | | | 48 | 25 | 3 |
| *EH108 1* | 0 | 3 | 21 | 45 | 35 | 5 |
| *EH108 2* | 0 | 5 | 25 | 55 | 22 | 4 |
| *EH111 1* | 0 | 0 | 45 | 55 | 32 | 4 |
| *EH111 2* | 0 | 1 | 52 | 62 | 35 | 3 |
| *EH111 3* | 0 | 0 | 40 | 45 | 31 | 11 |
| *MB235 A* | | | 15 | 47 | 62 | 41 |
| *MB235 B* | | | 19 | 64 | 60 | 38 |
| ***AVERAGE*** | **0** | **6** | **30** | **56** | **35** | **21** |
| ***STDEV*** | **0** | **8** | **13** | **12** | **16** | **18** |
| ***N*** | **7** | **7** | **9** | **20** | **20** | **20** |

**Supplemental Table 4. Flow Cytometry of Differentiation Time Course: PDGRα$^{+}$CD56$^{+}$%** (Data for Figure 1B)

| *Day* | *1* | *2* | *3* | *4* | *5* | *6* |
|---|---|---|---|---|---|---|
| *MB208 V1 31537_104918* | 0 | 2 | 10 | 34 | 73 | 83 |
| *MB208 V2 31537_105013* | 0 | 1 | 12 | 57 | 89 | 96 |
| *AAK11 1* | | | | 75 | 93 | 92 |
| *AAK11 2* | | | | 79 | 90 | 94 |
| *AAK11 3* | | | | 76 | 90 | 93 |
| *MB204 1* | | | | 66 | 86 | 80 |
| *MB204 2* | | | | 77 | 91 | 80 |
| *MB204 3* | | | | 71 | 89 | 80 |
| *MB204 4* | | | | 71 | 89 | 80 |
| *MB204 5* | | | | 66 | 84 | 80 |
| *MB204 6* | | | | 67 | 90 | 80 |
| *EH94* | | | | | | |
| *EH95* | | | | | | |
| *EH108 1* | 0 | 0 | 3 | 37 | 58 | 80 |
| *EH108 2* | 0 | 0 | 4 | 33 | 62 | 85 |
| *EH111 1* | 0 | 0 | 35 | 69 | 88 | 92 |
| *EH111 2* | 0 | 0 | 31 | 65 | 85 | 95 |
| *EH111 3* | 0 | 0 | 23 | 41 | 72 | 90 |
| *MB235 A* | | | 10 | 60 | 91 | 91 |
| *MB235 B* | | | 10 | 69 | 91 | 88 |
| ***AVERAGE*** | **0** | **0** | **15** | **62** | **84** | **87** |
| ***STDEV*** | **0** | **1** | **12** | **15** | **10** | **6** |
| ***N*** | **7** | **7** | **9** | **18** | **18** | **18** |

**Supplemental Table 5. Perforated Patch Clamp Action Potential Analysis** (Data for Figure 2E)

| *Cell* | MDP (mV) | APA (mV) | dV/dt max (V/s) | APD 20 (ms) | APD 50 (ms) | APD 90 (ms) | Cm (pF) |
|---|---|---|---|---|---|---|---|
| *24d13002_1Hz_24d13000* | -73.9 | 113.2 | 192.7 | 65.6 | 120.2 | 168.2 | 73.5 |
| *24d13005_1Hz_24d13005* | -66.3 | 76.3 | 163.3 | 14.9 | 35.9 | 63.5 | 107.9 |
| *24d13012_1Hz_24d13010* | -75.5 | 117.3 | 165.4 | 88.2 | 144.2 | 182.6 | 130 |
| *24d13019_1Hz_24d13017* | -60.1 | 99.8 | 27.7 | 42.7 | 70.3 | 90.1 | 127.6 |
| *24d13029_1Hz_24d13026* | -74.5 | 115.6 | 159.2 | 95.8 | 167.4 | 216.9 | |
| *24d13032_1Hz_24d13031* | -70.4 | 109.8 | 193.3 | 67.7 | 121.6 | 165.6 | 154.3 |
| *24d13041_1Hz_24d13039* | -73.1 | 116.3 | 72.0 | 105.5 | 167.1 | 205.2 | 165.8 |
| *24d13055_1Hz_24d13051* | -73.3 | 106.3 | 35.6 | 46.1 | 71.9 | 98.3 | 78.7 |
| *24n19002_1Hz_24n19001* | -72.2 | 108.2 | 24.8 | 29.1 | 51.5 | 87.5 | |
| *24n19013_1Hz_24n19010* | -81.3 | 99.5 | 36.7 | 42.8 | 67.3 | 91.9 | 79.3 |
| *24n19017_1Hz_24n19015* | -72.3 | 104.8 | 182.3 | 53.8 | 87.5 | 111.2 | 96 |
| *24n19025_1Hz_24n19024* | -67.7 | 104.6 | 17.1 | 93.1 | 152.3 | 177.0 | |
| *24n19028_1Hz_24n19026* | -78.0 | 116.8 | 20.6 | 82.3 | 150.2 | 226.7 | |
| *24n19033_1Hz_24n19031* | -69.0 | 111.3 | 115.7 | 95.2 | 166.5 | 207.8 | 95.4 |
| *24n19039_1Hz_24n19038* | -68.1 | 112.0 | 164.5 | 86.2 | 139.3 | 166.8 | |
| *24n19041_1Hz_24n19040* | -68.5 | 106.0 | 31.1 | 61.5 | 129.3 | 179.8 | |
| *24n19044_1Hz_24n19043* | -72.4 | 100.3 | 12.9 | 63.0 | 134.0 | 182.3 | |
| *24n19051_1Hz_24n19050* | -73.4 | 102.3 | 141.8 | 31.3 | 53.9 | 76.9 | 69.6 |
| *24n19056_1Hz_24n19053* | -88.7 | 124.3 | 22.3 | 39.0 | 81.0 | 341.2 | 92.1 |
| *24n22001_1Hz_24n22000* | -69.3 | 97.5 | 149.7 | 38.6 | 66.5 | 90.9 | 96.7 |
| *24n22007_1Hz_24n22005* | -72.1 | 103.5 | 263.3 | 37.9 | 69.2 | 100.6 | 90.7 |
| *24n22011_1Hz_24n22009* | -65.5 | 102.3 | 87.5 | 71.2 | 119.0 | 145.8 | |
| *24n22016_1Hz_24n22014* | -72.2 | 106.5 | 208.4 | 45.6 | 88.3 | 135.2 | 116.4 |
| *24n22021_1Hz_24n22020* | -70.0 | 108.1 | 156.0 | 62.4 | 96.4 | 126.4 | 124 |
| *24n22025_1Hz_24n22024* | -72.4 | 109.7 | 213.5 | 80.2 | 119.3 | 144.3 | |
| *24n22031_1Hz_24n22028* | -77.7 | 107.6 | 59.5 | 46.0 | 68.8 | 88.8 | 73.3 |
| ***Average*** | **-72.5** | **107.0** | **112.1** | **61.9** | **106.8** | **150.3** | **108.5** |
| ***SD*** | **5.50** | **8.81** | **75.50** | **24.23** | **39.99** | **61.30** | **33.14** |
| ***SEM*** | **1.06** | **1.70** | **14.53** | **4.66** | **7.70** | **11.80** | **7.81** |
| ***N*** | **27.0** | **27** | **27** | **27** | **27** | **27** | **18** |

**Supplemental Table 6. Summary of Randomized Pig Assignments and Studies**

| Pig ID | Sex | Weight (kg) | Treatment | MRI Endpoint at 4 wk | Loop Recorder | P-V Loop Analysis | Survival | Graft Detected (Ku80+) |
|---|---|---|---|---|---|---|---|---|
| 1423 | M | 30 | CCP | No* | Yes | No | 4-wk | Yes |
| 1425 | M | 30 | Control | No# | No# | No | 8-wk | N/A |
| 1429 | M | 34 | cECM | No* | Yes | No | 4-wk | N/A |
| 1432 | M | 32 | CCP+cECM | No* | Yes | No | 3-wk | Yes |
| 1437 | M | 28 | CCP | No* | Yes | No | 5-wk | Yes |
| 1444 | M | 30 | Control | No^ | Yes | No | 5-wk | N/A |
| 1421 | M | 29 | CCP | No | Yes | No | 7-wk | Yes |
| 1422 | M | 30 | CCP+cECM | No* | Yes | No | 7-wk | Yes |
| 1424 | M | 31 | Control | No* | Yes | No | 5-wk | N/A |
| 1434 | M | 31 | CCP | No* | Yes | No | 5-wk | Yes |
| 1436 | M | 30 | CCP+cECM | No* | Yes | No | 4-wk | No |
| 2304 | F | 30 | cECM | No# | No | No | 8-wk | N/A |
| 2306 | F | 31 | cECM | Yes | Yes | No | 6-wk | N/A |
| 2310 | F | 32 | CCP | Yes | No | Yes | 8-wk | Yes |
| 2322 | M | 32 | Control | No* | No | No | 5-wk | N/A |
| 2339 | M | 30 | CCP+cECM | Yes | Yes | Yes | 8-wk | Yes |
| 2235 | M | 30 | cECM | Yes | Yes | No | 4-wk | N/A |
| 2283 | M | 32 | Control | Yes | Yes | Yes | 7-wk | N/A |
| 2288 | M | 32 | cECM | Yes | Yes | Yes | 8-wk | N/A |
| 2300 | F | 30 | CCP+cECM | No# | Yes | Yes | 8-wk | Yes |
| 2302 | F | 31 | CCP | No* | Yes | No | 3-wk | No |
| 2303 | F | 30 | CCP+cECM | Yes | No# | No | 7-wk | Yes |
| 2990 | F | 32 | CCP | Yes | No# | Yes | 4-wk | Yes |
| 3153 | M | 33 | Control | Yes | No# | Yes | 4-wk | N/A |
| 3162 | M | 35 | CCP | Yes | No# | Yes | 4-wk | Yes |
| 3181 | F | 33 | CCP+cECM | Yes | No# | Yes | 4-wk | Yes |
| 3702 | M | 29 | Control | Yes | Yes | Yes | 4-wk | N/A |
| 3736 | F | 34 | Control | Yes | Yes | Yes | 4-wk | N/A |
| 3708 | M | 30 | CCP | Yes | Yes | Yes | 4-wk | Yes |
| 3726 | F | 34 | CCP+cECM | Yes | Yes | Yes | 4-wk | Yes |
| 3720 | F | 29 | CCP | No* | Yes | No | 4-wk | Yes |
| 3996 | M | 30 | CCP+cECM | No* | Yes | No | 4-wk | Yes |
| 4045 | M | 30 | Control | Yes | Yes | Yes | 4-wk | N/A |
| 4309 | M | 30 | CCP+cECM | Yes | Yes | Yes | 4-wk | Yes |
| 4433 | M | 32 | CCP | Yes | Yes | Yes | 4-wk | Yes |
| 4472 | F | 31 | Control | Yes | Yes | Yes | 4-wk | N/A |
| 4680 | F | 31 | CCP+cECM | Yes | Yes | Yes | 4-wk | Yes |
| 504 | M | 40 | CCP+cECM | Yes | Yes | Yes | 4-wk | Yes |
| 507 | M | 40 | CCP | No# | Yes | Yes | 4-wk | Yes |
| 582 | F | 28 | Control | No* | Yes | No | 3-wk | N/A |
| 598 | M | 28 | CCP | No* | Yes | No | 4-wk | Yes |
| 650 | F | 30 | CCP | No* | Yes | No | 3-wk | Yes |
| 654 | F | 30 | Control | Yes | Yes | Yes | 4-wk | N/A |
| 672 | M | 30 | CCP+cECM | Yes | Yes | Yes | 4-wk | Yes |
| 621 | M | 31 | Control | Yes | Yes | Yes | 4-wk | N/A |
| 622 | M | 29 | CCP+cECM | Yes | Yes | Yes | 4-wk | Yes |

Total Randomized pigs:46
*Severe infection
#Technical failure
^Infarct<8%, EF>45%

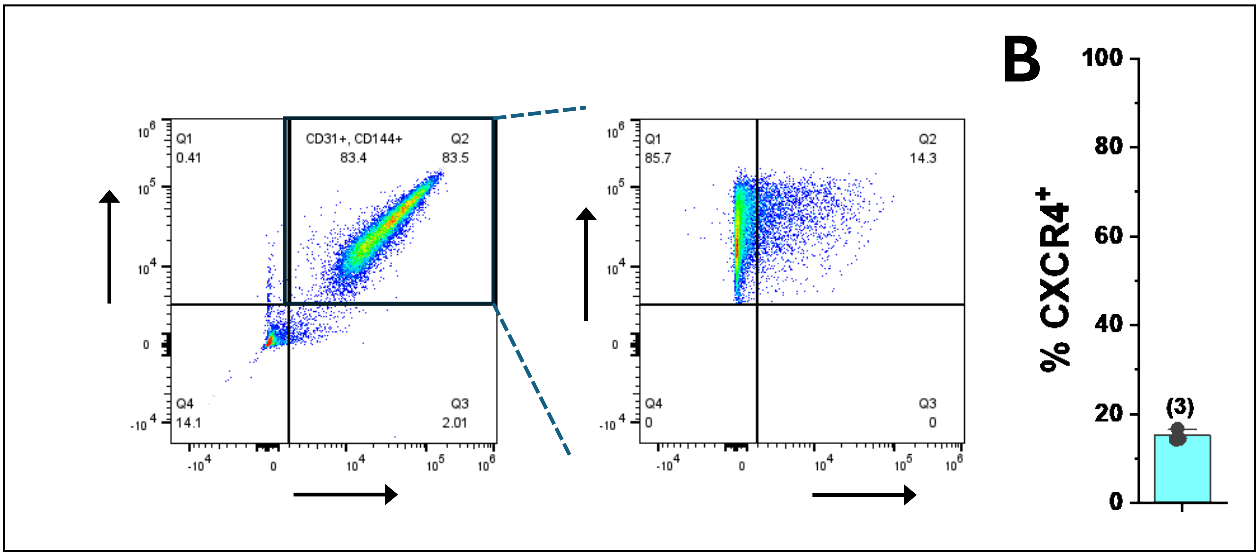


**Figure S1. CCP differentiated iPSC-EC characterization. A)** Representative flow cytometry after 12 days culture of CCPs in Vasculife LS1020$^{TM}$ medium with sequential gating for endothelial markers, CD31 and CD144, followed by gating of the double positive population for CXCR4, a marker for arterial ECs. **B)** Plot of percentage of CD31$^+$/CD144$^+$ cells that positive for the CXCR4 with bar being average and error bar representing SD.

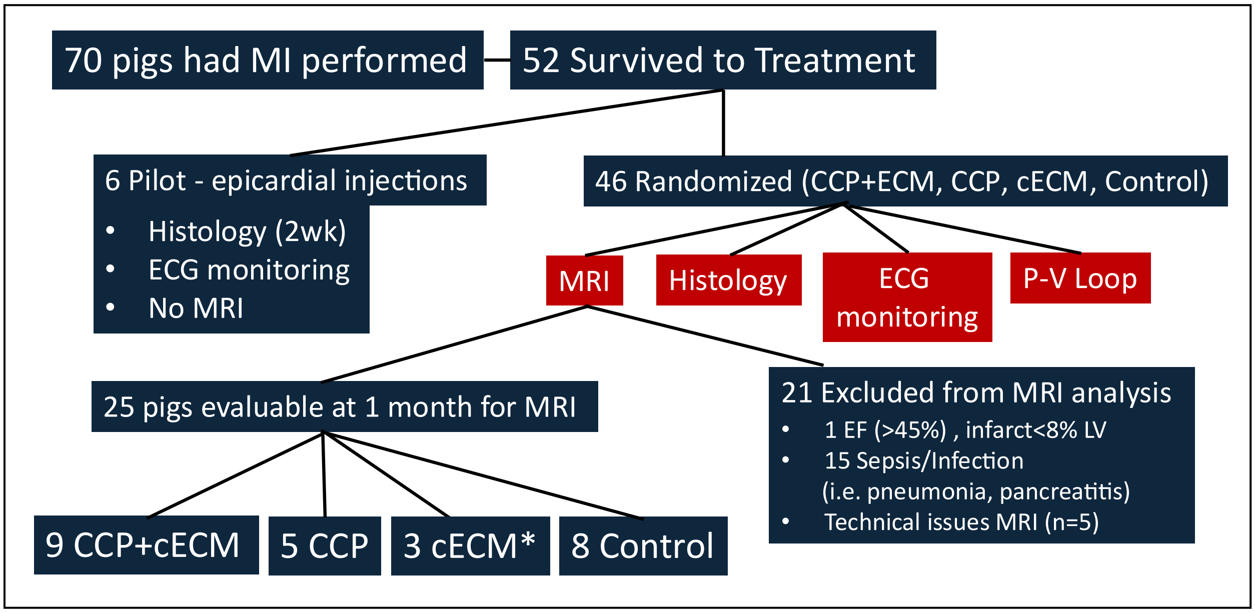


**Figure S2. Allocation of pigs**. Flowchart showing pig allocation following the MI to an initial pilot feasibility cohort and to the 4-group randomized cohort with reasons for exclusions from the final analysis. *cECM cohort stopped early to prioritize other groups.

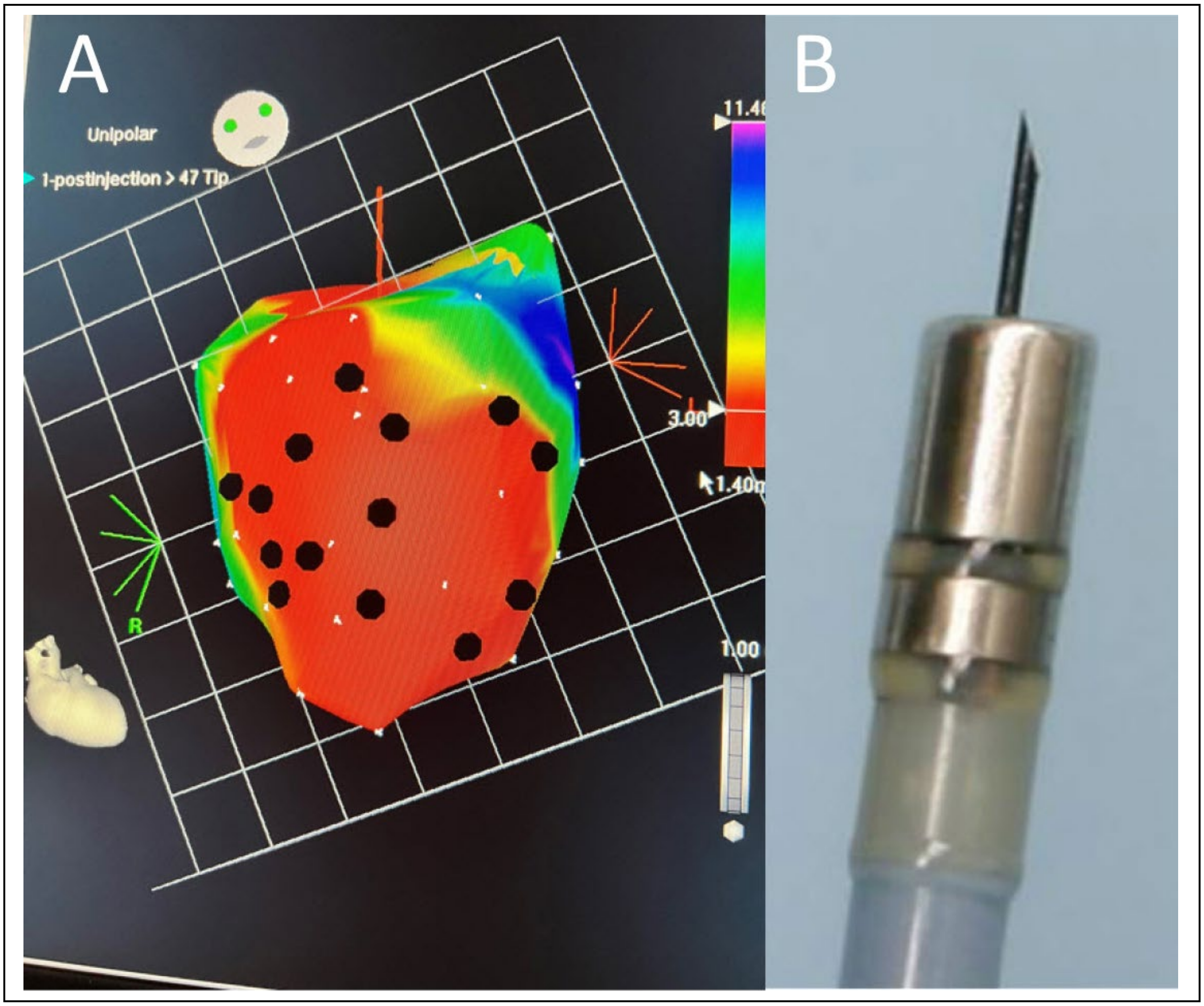


**Figure S3**. **Electroanatomic mapping and cell delivery. A)** Representative image (pig 672) of a cardiac unipolar voltage map showing the large infarct zone (red) with 14 annotated injection locations distributed within the infarct and in proximity to the peri-infarct border zone. **B)** The Myostar, straight needle tipped catheter that is trackable on the electroanatomical mapping system, was used to perform the injections.

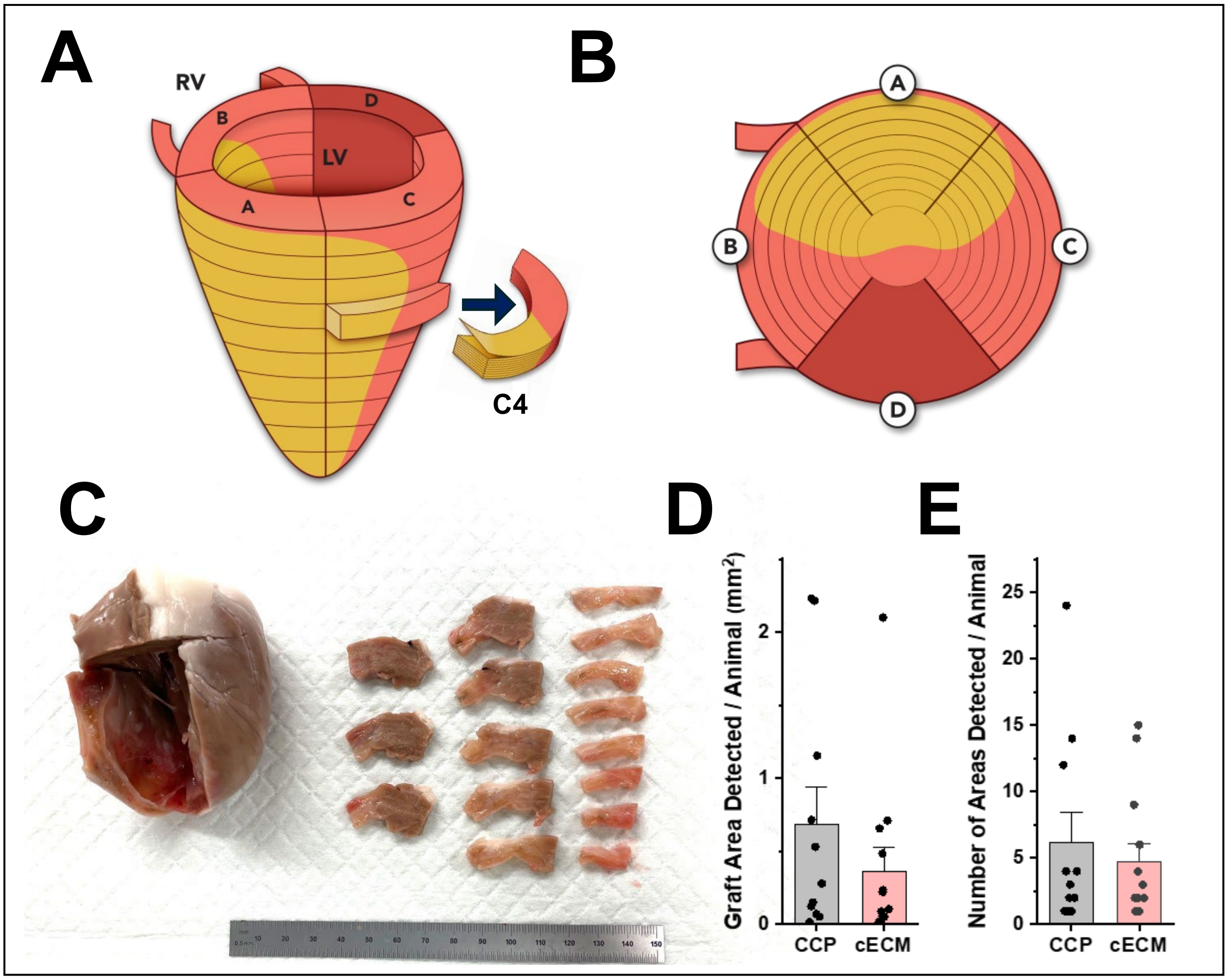


**Figure S4. Blocking strategy for histopathology assessment of porcine hearts. A)** Schematic diagram for blocking left ventricle from area above infarct zone to the apex. The ventricle is divided into 4 zones, A (anterior wall encompassing most of infarct), B (septum with infarct and border zone), C (anterolateral wall with infarct and borderzone), and D (remote zone). Zones A, B, and C are then dissected into blocks of 1.0-1.5 mm, fixed and paraffin embedded for histological assessment as described in Methods. Blocks are uniquely named and tracked with an example of the C4 block shown. **B)** Bullseye plot schematic of left ventricle showing a polar map of 4 regions and areas blocked for histology. **C)** Gross porcine heart in which region A (8 left tissue blocks) and region C (8 right most tissue blocks) have been dissected. **D)** Total graft area detected in screened sections per animal. **E)** Number of areas detected in screened sections per animal. Bars indicate mean and error bars are S.E.M.

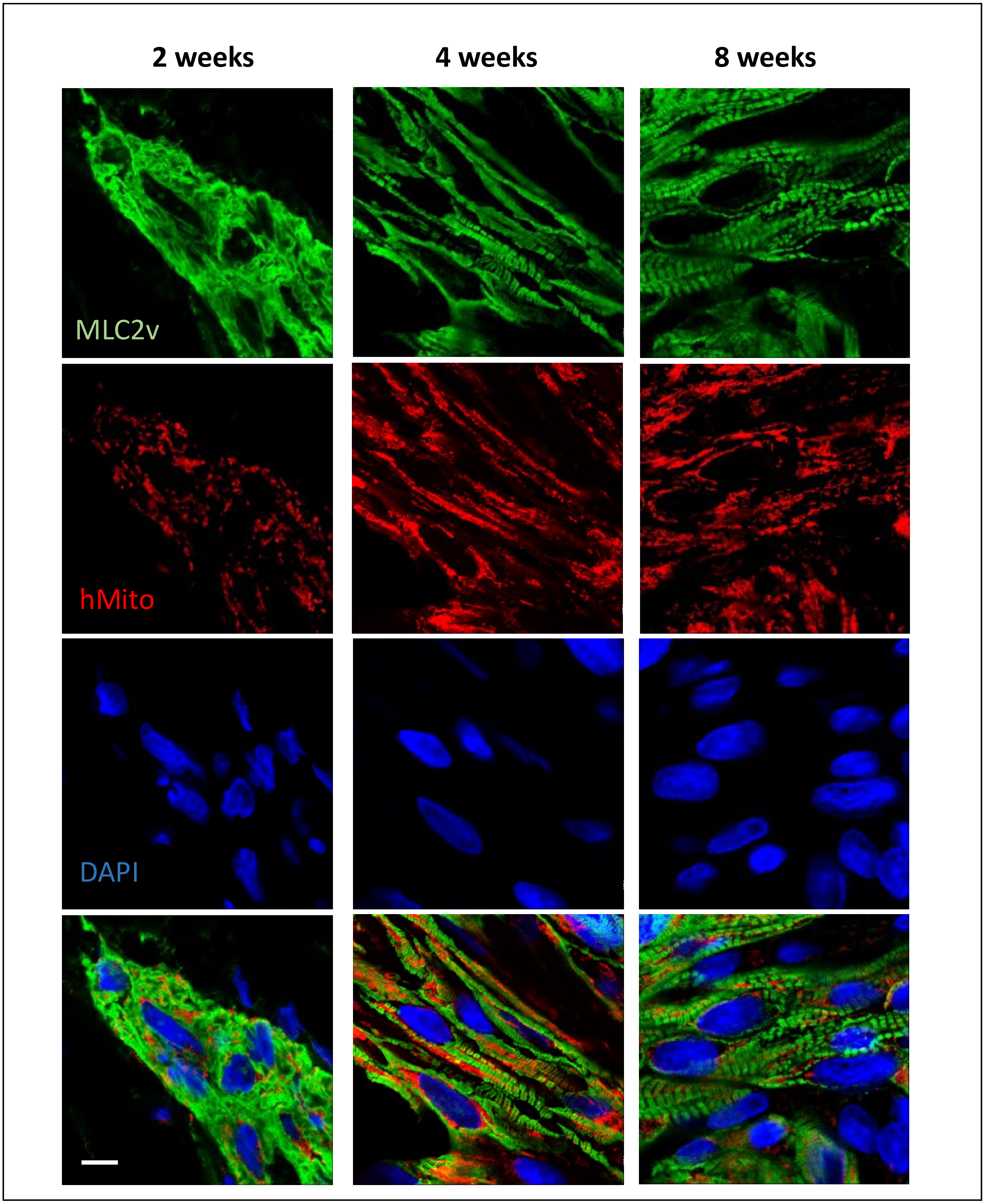


**Figure S5.** Confocal immunofluorescence imaging showing CCP-derived grafts at time points indicated post-delivery labeling with ventricular-specific MLC2v, human mitochondrial antibody and DAPI shown in each channel and overlaid in bottom row which correspond to Figure 4 J,K,L.
